# Manganese transport by *Streptococcus sanguinis* in acidic conditions and its impact on growth *in vitro* and *in vivo*

**DOI:** 10.1101/2021.05.28.446192

**Authors:** Tanya Puccio, Alexander C. Schultz, Claudia A. Lizarraga, Ashley S. Bryant, David J. Culp, Robert A. Burne, Todd Kitten

## Abstract

*Streptococcus sanguinis* is an oral commensal and an etiological agent of infective endocarditis. Previous studies have identified the SsaACB manganese transporter as essential for endocarditis virulence; however, the significance of SsaACB in the oral environment has never been examined. Here we report that a Δ*ssaACB* mutant of strain SK36 exhibits reduced growth and manganese uptake under acidic conditions. Further studies revealed that these deficits resulted from the decreased activity of TmpA, shown in the accompanying paper to function as a ZIP-family manganese transporter. Transcriptomic analysis of fermentor-grown cultures of SK36 WT and Δ*ssaACB* strains identified pH-dependent changes related to carbon catabolite repression in both strains, though their magnitude was generally greater in the mutant. In strain VMC66, which possesses a MntH transporter, loss of SsaACB did not significantly alter growth or cellular manganese levels under the same conditions. Interestingly, there were only modest differences between SK36 and its Δ*ssaACB* mutant in competition with *Streptococcus mutans in vitro* and in a murine oral colonization model. Our results suggest that the heterogeneity of the oral environment may provide a rationale for the variety of manganese transporters found in *S. sanguinis* and point to strategies for enhancing the safety of oral probiotics.

**Graphical Abstract:** Depiction of methods. Streptococcal strains used are depicted at the top. The four methods illustrated are: 1. *S. sanguinis* cells were grown in media at pH 7.3 and pH 6.2 and quantified by plating. 2. *S. sanguinis* cells were grown in a fermentor vessel for RNA-sequencing and metal analysis. 3. *S. sanguinis* and *S. mutans* cells were grown in 12-well plates singly or in competition, then plated and the pH of the media measured. 4. *S. sanguinis* and *S. mutans* cells were inoculated into the mouths of mice. Oral swabs and dental biofilms from molars were assayed for bacterial composition by qPCR. (Biorender)

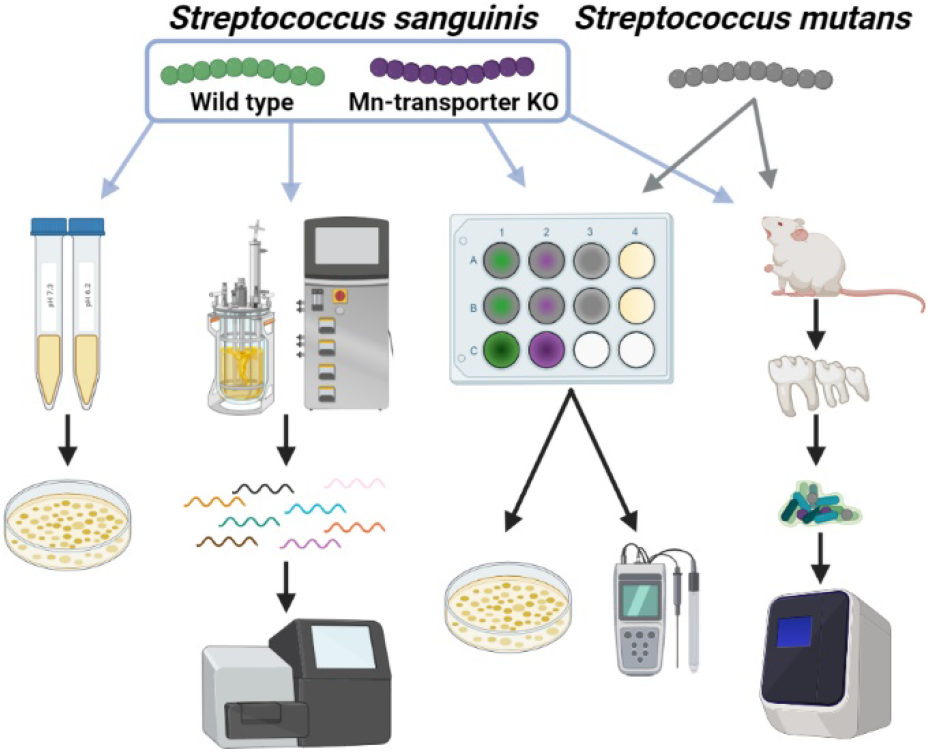

## Introduction

The oral cavity is a complex environment with dynamic microbial communities, fluctuating metabolite availability, and diverse habitats (Lamont *et al.*, 2018, Aas *et al.*, 2005, Marsh *et al.*, 2016, Kolenbrander, 2011, Jakubovics, 2015, Kolenbrander *et al.*, 2010, Chen *et al.*, 2010). Resident microorganisms must compete for space and nutrients within each of these habitats, including the dental biofilm known as plaque. Some species, such as *Streptococcus mutans*, utilize acid production to carve out their own niche by lowering the pH of the local environment. This allows competition with other less aciduric (acid-tolerant) species but also results in demineralization of the tooth enamel, or dental caries (Gross *et al.*, 2012, Nascimento *et al.*, 2009, Moye *et al.*, 2014).

One of the species *S. mutans* competes with is *Streptococcus sanguinis*. While *S. sanguinis* is acidogenic (acid-producing), it lacks the aciduricity of *S. mutans* (Bender *et al.*, 1986, Diaz-Garrido *et al.*, 2020, Sasaki *et al.*, 2018, Svensater *et al.*, 1997). Conversely, *S. sanguinis* possesses the ability to produce (Kreth *et al.*, 2008, Garcia-Mendoza *et al.*, 1993) and survive in (Xu *et al.*, 2014) large quantities of hydrogen peroxide (H_2_O_2_), which is thought to enhance its ability to compete against *S. mutans*, which does not produce H_2_O_2_ (Kreth *et al.*, 2005). Thus, *S. sanguinis* is typically found in much greater abundance at healthy oral sites while *S. mutans* predominates in carious lesions (Giacaman *et al.*, 2015, Belda-Ferre *et al.*, 2012, Gross *et al.*, 2012).

Oral bacteria possess a variety of mechanisms for tolerating acid stress (Papadimitriou *et al.*, 2016, Guan & Liu, 2020, Liu *et al.*, 2015, Quivey *et al.*, 2001, Cotter & Hill, 2003). The *S. sanguinis* genome (Xu *et al.*, 2007) encodes multiple systems that are predicted to play a role: F_0_F_1_-ATPases (Kuhnert & Quivey, 2003); an arginine deiminase system (Burne *et al.*, 1989, Curran *et al.*, 1995, Floderus *et al.*, 1990); various chaperones and proteases (Shabayek & Spellerberg, 2017, Lemos *et al.*, 2001, Lemos & Burne, 2002); and superoxide dismutase (Kim *et al.*, 2005, Wen & Burne, 2004, Crump *et al.*, 2014). *S. sanguinis* also appears to possess an acid tolerance response (ATR), whereby brief exposure to sub-lethal acid levels protects against a lethal drop in pH (Cotter & Hill, 2003), although it is less effective than that in the related species *Streptococcus gordonii* (Cheng *et al.*, 2018). Each of these systems likely contributes to the survival of *S. sanguinis* in acidic environments.

Manganese (Mn) has recently been implicated in acid stress tolerance in *Streptococcus agalactiae* (Shabayek *et al.*, 2016) and *S. mutans* (Kajfasz *et al.*, 2020). In both species, loss of a natural resistance-associated macrophage protein (NRAMP) family manganese transporter, MntH, led to a reduction in growth in acidic conditions. In *S. mutans*, the loss of the lipoprotein component of the ABC manganese transporter, SloC, did not impact growth in acidic media unless the *sloC* mutation was combined with the *mntH* mutation (Kajfasz *et al.*, 2020). Similarly, it was previously determined that the *fimCBA* operon was not important for acid tolerance in *Streptococcus parasanguinis* (Chen *et al.*, 2013), which also possesses a *mntH* gene. Manganese transport has also been examined in *S. sanguinis*, which possesses an analogous ABC transport system encoded by the *ssaACB* operon but not a *mntH* gene in most strains. These studies have established the importance of manganese not for acid tolerance, but for virulence in relation to the disease infective endocarditis (Crump *et al.*, 2014, Baker *et al.*, 2019). *S. sanguinis* is a frequent cause of this serious illness, which exhibits a 12-30% mortality rate (Bor *et al.*, 2013, Cahill *et al.*, 2017, Jamil *et al.*, 2019, Wilson *et al.*, 2021) and for which the only form of prevention—antibiotic prophylaxis—is controversial. This has led to the investigation of SsaB, the lipoprotein component of the ABC manganese transport system, as a target for endocarditis prevention. Given the association of *S. sanguinis* with health in the oral cavity, targeting of SsaB would be of greater value if this does not compromise oral competitiveness.

Here we report for the first time the growth of a naturally MntH-deficient streptococcal strain, *S. sanguinis* SK36, under reduced-pH conditions and for the first time in any bacterium, the effect of reduced pH on manganese transport by a newly described ZIP family transporter. We then assess the growth of a MntH-encoding strain of *S. sanguinis*, VMC66, in reduced-pH conditions. Finally, we report the effect of deleting the ABC manganese transporter on the competitive fitness of *S. sanguinis* against *S. mutans in vitro* and *in vivo*.

## Results

### Growth of *S. sanguinis* at reduced pH

As a first step in determining whether manganese transport is important for growth at reduced pH in S*. sanguinis*, we employed a mutant of SK36 in which the ABC manganese transporter SsaACB has been deleted (Baker *et al.*, 2019). We assessed the growth of wild-type (WT) SK36 and its Δ*ssaACB* single transport-system mutant in brain heart infusion (BHI) broth at its unadjusted pH (~7.3) and under increasingly acidic conditions (Figure 1A). We have recently demonstrated that this mutant grows significantly less well than WT in a low-manganese medium, serum, in 6% O_2_ but indistinguishably in 1% O_2_ (Puccio *et al.*, 2021a). To rule out oxidative stress as a confounding factor, cells were grown in microaerobic conditions (1% O_2_). At pH 7.3, the culture densities of the two strains were not significantly different from each other. When the pH was lowered to 6.4 and 6.3, the final density of the WT culture was not significantly different from that at pH 7.3 whereas the final density of the Δ*ssaACB* mutant progressively decreased. The final density of both cultures at pH 6.2 was significantly less than at pH 7.3, although the difference was much greater for the Δ*ssaACB* mutant than for WT.

**Figure 1.**
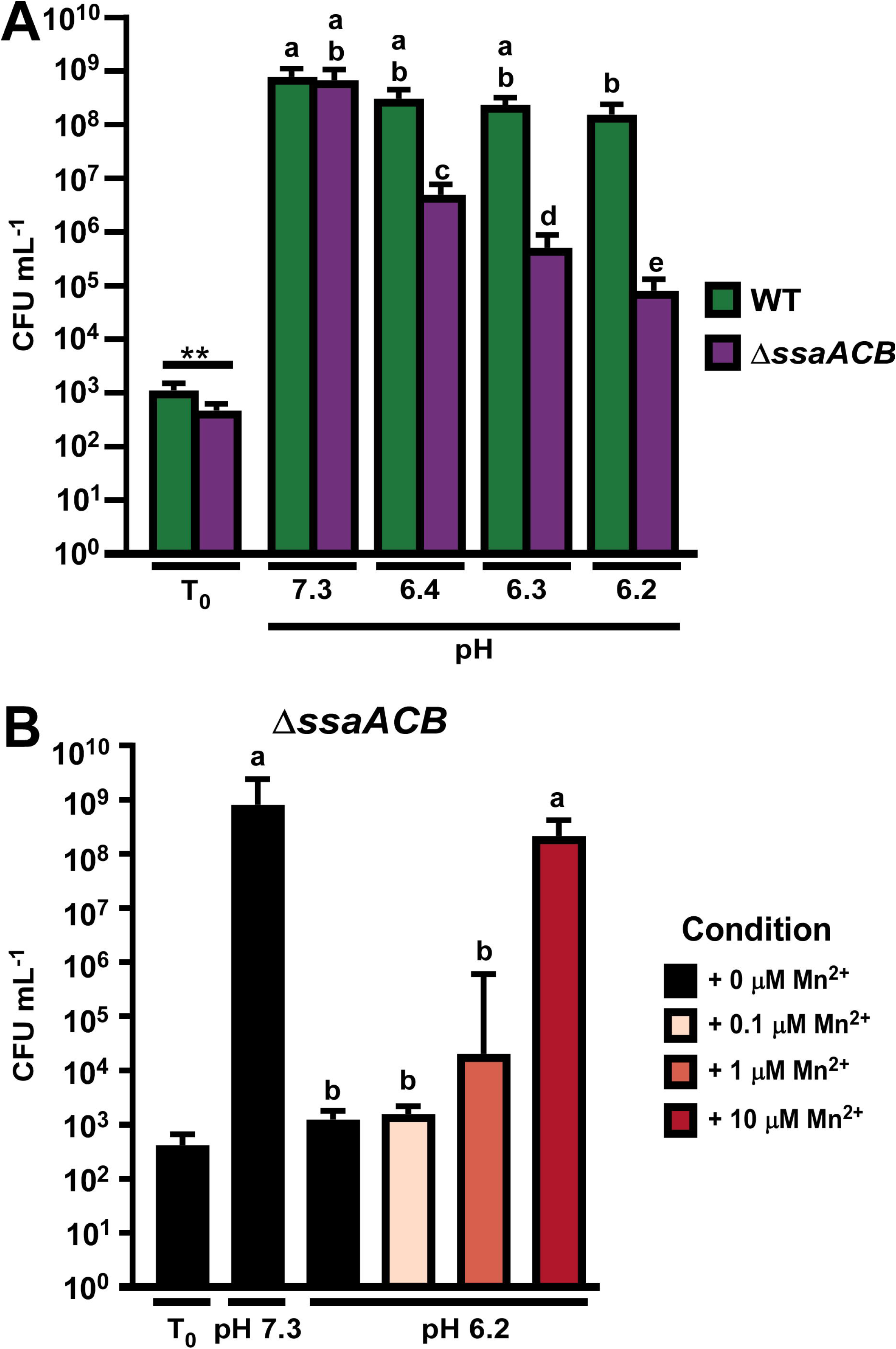
Growth of the *S. sanguinis* SK36 WT and an Δ*ssaACB* mutant in reduced-pH BHI. BHI at different pH levels was preincubated at 1% O_2_ and inoculated from an overnight culture. Cultures were incubated for 24 h prior to plating. (A) Growth in BHI at various pH levels was assessed. (B) Exogenous Mn^2+^ was added at listed concentrations. Means and standard deviations of at least three replicates are displayed. Significance was determined by unpaired two-tailed t-tests for T_0_ values and one-way ANOVA with a Tukey multiple comparisons post-test for T_24_ values. Bars with the same letter are not significantly different from each other (*P* > 0.05).

Although we suspected that the growth arrest of the Δ*ssaACB* mutant at pH 6.2 was due to lower manganese levels in this strain (Murgas *et al.*, 2020), we wanted to test this hypothesis directly. We previously measured the concentration of manganese in BHI from the same supplier as that used in the present study by inductively coupled plasma mass spectrometry (ICP-MS) and found that it averaged 0.36 μM (Murgas *et al.*, 2020). We added exogenous Mn^2+^ to pH 6.2 cultures of the Δ*ssaACB* mutant and found that the addition of 10 μM Mn^2+^ was sufficient to rescue the growth (Figure 1B), suggesting that the reduced manganese levels found in this mutant (Murgas *et al.*, 2020) were contributing to the reduced ability to tolerate low pH conditions.

### Contribution of secondary manganese transporters to acid tolerance

The success of the manganese rescue experiment shown in Figure 1B suggests the existence of an additional mechanism for manganese uptake, albeit one with lower affinity than the SsaACB transporter. In the accompanying manuscript (Puccio *et al.*, 2021a), we report the identification and characterization of a ZIP family protein, TmpA, as a secondary manganese transporter present in at least 64 *S. sanguinis* strains whose sequences are available in the GenBank whole-genome shotgun contigs database. Additionally, we noted the presence of a gene encoding a third manganese transporter from the NRAMP family (Nevo & Nelson, 2006), MntH, in eight of these strains, one of which was VMC66. We hypothesized that each of the three *S. sanguinis* manganese transporters are at least partly redundant but may have optimal activity under differing conditions. In *S. agalactiae*, MntH functions best at acidic pH and its gene is expressed highly at pH 5 (Shabayek *et al.*, 2016). The ZIP-family protein in *Escherichia coli* was found to function best near neutral pH (Taudte & Grass, 2010) and we have yet to determine a condition where the *tmpA* gene in *S. sanguinis* is differentially expressed (Puccio *et al.*, 2021a).

To determine whether TmpA contributes to the growth of SK36 WT or Δ*ssaACB* mutant cells at normal or reduced pH, we assessed the growth of Δ*tmpA* mutants in both backgrounds in BHI pH 7.3 and 6.2 in an atmosphere of 1% O_2_ (Figure 2A). Growth of the Δ*tmpA* mutant was not affected at either pH, indicating that SsaACB transports sufficient manganese at pH 6.2 to maintain growth. The double Δ*ssaACB* Δ*tmpA* mutant grew poorly at both pH 7.3 and 6.2 and its growth was not significantly different from that of its Δ*ssaACB* parent at pH 6.2.

**Figure 2.**
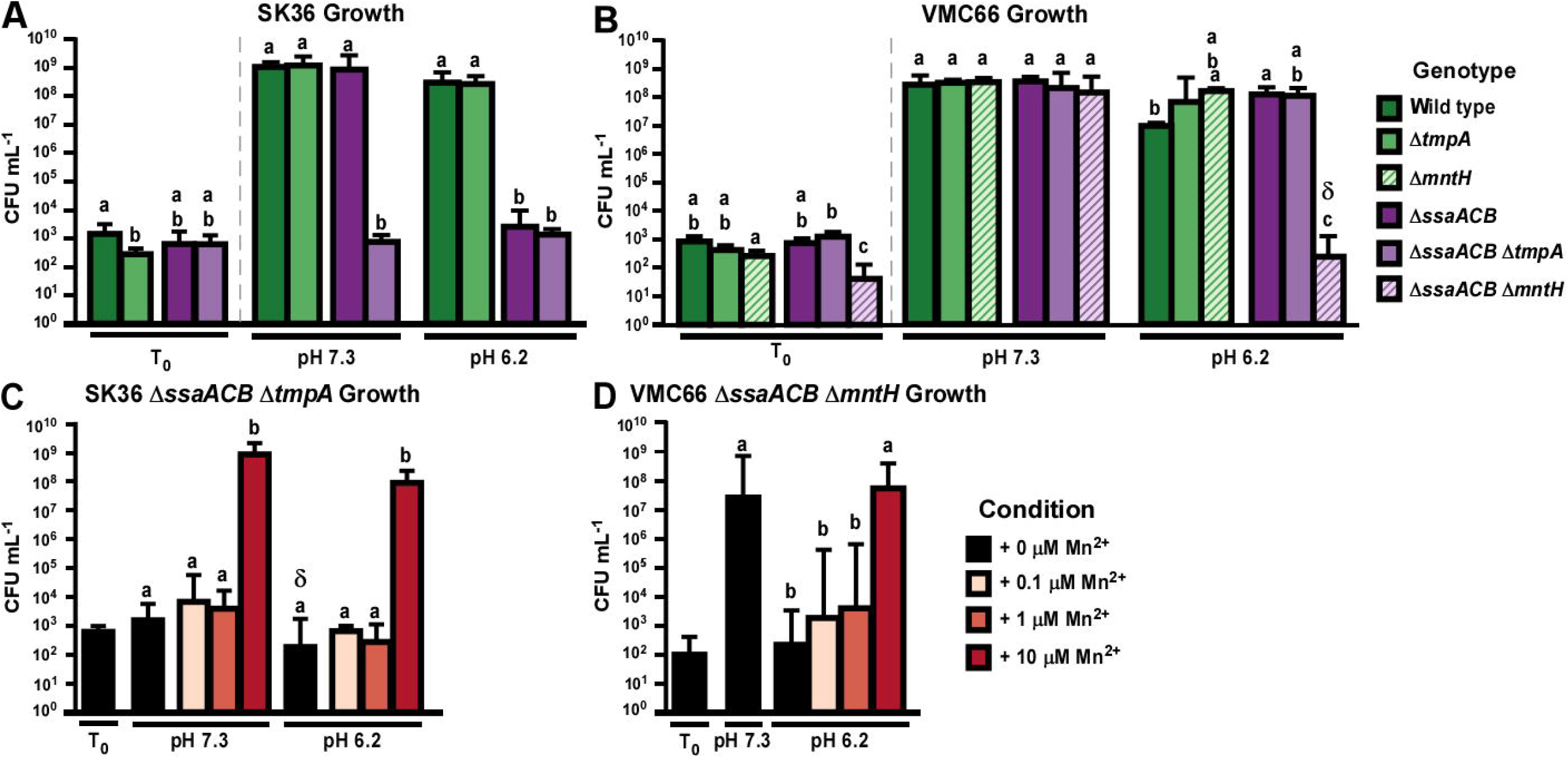
Effect of the loss of manganese transporters on growth of *S. sanguinis* SK36 and VMC66 strains in reduced-pH BHI. Growth of manganese transporter mutants in the SK36 (A) and VMC66 (B) backgrounds was assessed in BHI at pH 7.3 and pH 6.2 in 1% O_2_. Strains that grew poorly in A & B were assessed for growth under the same conditions ± various concentrations of Mn^2+^ (C-D). Means and standard deviations of at least three replicates are displayed. Significance was determined by one-way ANOVA with a Tukey multiple comparisons post-test separately for T_0_ and T_24_ values (vertical dashed lines). Bars with the same letter within a chart are not significantly different (*P* > 0.05). Bars with δ indicate that at least one replicate fell below the limit of detection.

In the VMC66 background, we generated mutants deleted for each of the transporters: Δ*tmpA*, Δ*mntH*, and Δ*ssaACB*, as well as double mutants lacking SsaACB and one of the secondary transporters: Δ*ssaACB* Δ*tmpA*, and Δ*ssaACB* Δ*mntH.* We then assessed their growth in BHI pH 7.3 and pH 6.2 at 1% O_2_ (Figure 2B). All strains grew identically at pH 7.3 but growth of the Δ*ssaACB* Δ*mntH* strain was dramatically reduced at pH 6.2.

We also wanted to assess whether the decreased growth of the SK36 Δ*ssaACB* Δ*tmpA* mutant and the VMC66 Δ*ssaACB* Δ*mntH* mutant observed in Figure 2A-B was primarily due to decreased manganese levels in these mutants. Growth of both mutants was significantly improved by the addition of 10 μM Mn^2+^ (Figure 2C-D).

We next wanted to determine how the reduced pH affected manganese levels in these strains. In our accompanying study (Puccio *et al.*, 2021a), we determined the average manganese levels of the SK36 strains in BHI pH 7.3 using inductively coupled plasma optical emission spectroscopy (ICP-OES), which are reproduced in Figure 3A. The mean levels of manganese were lower in these strains in BHI at pH 6.2 (Figure 3B); so much so that levels in both Δ*ssaACB* and Δ*ssaACB* Δ*tmpA* fell below the limit of detection unless exogenous Mn^2+^ was added. While there was a significant difference between the Δ*ssaACB* Δ*tmpA* mutant and its parent when 10 μM Mn^2+^ was added at pH 7.3 (Figure 3A), there were no significant differences between either parent strain (WT or Δ*ssaACB*) and its corresponding Δ*tmpA* mutant in any condition at pH 6.2 (Figure 3B). Levels of the biologically relevant metals iron (Fe), zinc (Zn), and magnesium (Mg) were not significantly affected by the lack of *tmpA* in either the WT or Δ*ssaACB* background in BHI at pH 7.3 (Puccio *et al.*, 2021a) or pH 6.2 ± 10 μM Mn^2+^ (Figure S1A, C, E).

**Figure 3.**
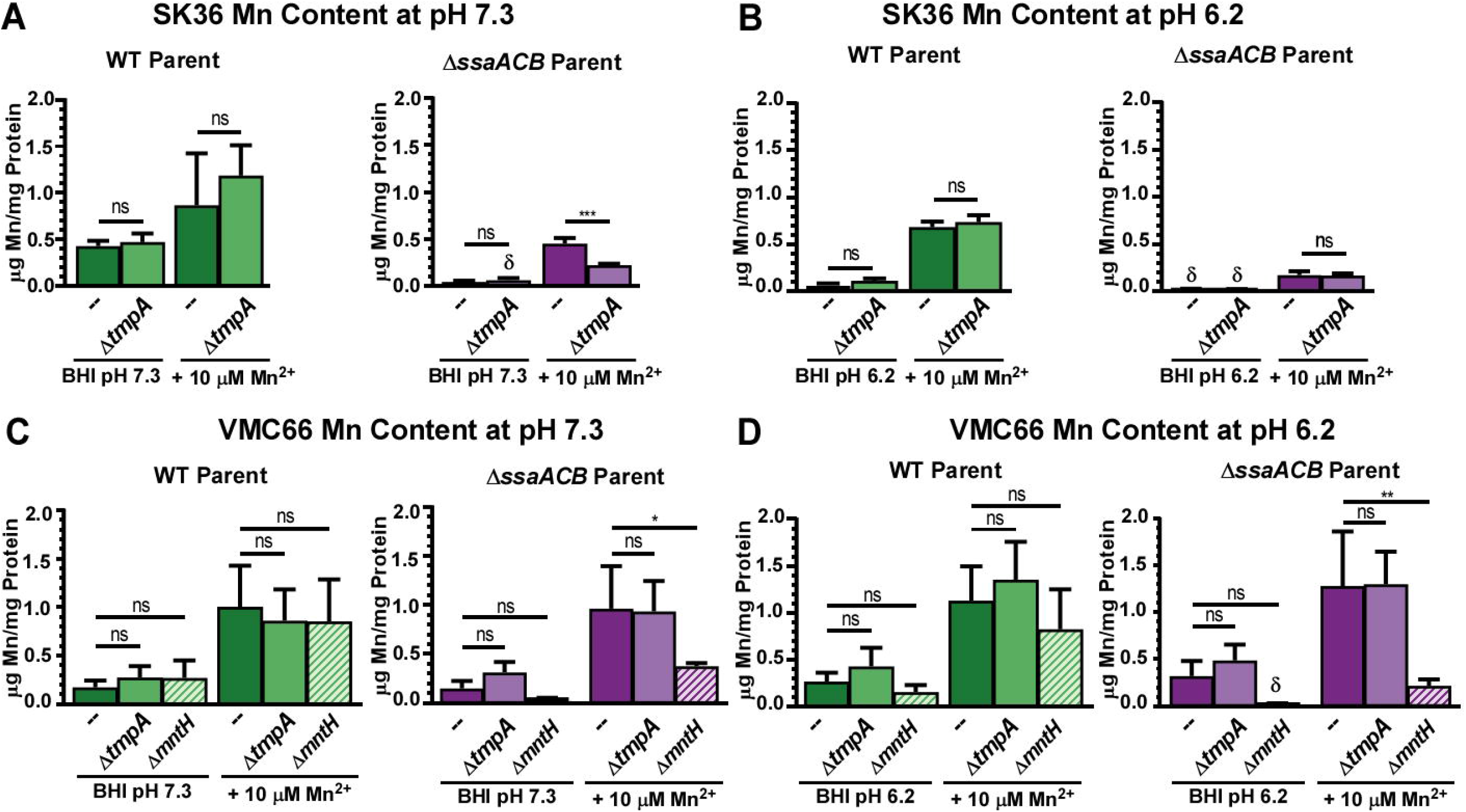
Manganese content of *S. sanguinis* SK36 and VMC66 and their respective manganese transporter mutants. Manganese content of transporter mutants in the SK36 (A & B) and VMC66 (C & D) backgrounds in BHI ± 10 μM Mn^2+^ at pH 7.3 (A-C) and pH 6.2 (B-D) was assessed by ICP-OES. Means and standard deviations of at least three replicates are displayed. Significance was determined by one-way ANOVA with Bonferonni’s multiple comparisons tests comparing each mutant strain to its respective parent under the same growth conditions. Bars indicated by δ had at least one value that fell below the lowest standard. Data used in A & B are also included in the accompanying paper (Puccio *et al.*, 2021a).

Manganese levels of VMC66 and derivative strains in BHI at pH 7.3 were also measured in Puccio *et al.* (2021a) and are shown in Figure 3C. With the exception of both Δ*mntH* strains, the mean manganese levels at pH 6.2 (Figure 3D) were higher than at pH 7.3. The loss of TmpA did not impact manganese levels in either the WT VMC66 or the Δ*ssaACB* mutant under either pH condition. Levels of iron, zinc, and magnesium were not significantly affected by deletion of the *tmpA* or *mntH* gene in either the WT or Δ*ssaACB* background at pH 7.3 (Puccio *et al.*, 2021a) or pH 6.2 (Figure S1B, D, F).

### Effect of pH reduction on SK36 WT and Δ*ssaACB* growth in a fermentor

To assess the effect of reduced pH on the transcriptome of SK36 WT and Δ*ssaACB* mutant cells, we employed the use of a fermentor (Figure S2). To minimize the impact of oxidative stress, we used the minimum possible airflow (0.03 lpm) throughout the experiment. We were unable to turn off the air entirely because cultures without airflow grew poorly (data not shown). Cells were grown at pH 7.4 (pH of human blood) until the OD peaked, and then for 1 hr with a media flow rate of 700 mL h^−1^ before collection of the T_−20_ min sample for RNA-seq. Twenty minutes later (T_0_), the pH was changed to 6.2 by addition of 2 N HCl and 2 N KOH in response to the output from an indwelling pH probe. A pH of 6.2 was chosen because it did not affect WT (Figure S2A) but led to a decrease in growth rate of the Δ*ssaACB* mutant as evidenced by the drop in OD (Figure S2B). Under constant-flow conditions, a drop in OD is expected when a culture’s doubling rate drops below its dilution rate (Burne & Chen, 1998). Samples at pH 6.2 were removed at T_10_, T_25_, and T_50_ min and RNA was isolated for RNA-seq analysis.

We next wanted to determine if manganese levels were affected by pH reduction in the fermentor. We assessed the metal content of cells at each sample time point using ICP-OES (Figure 4). Manganese levels significantly decreased in both strains, further confirming that reduced pH conditions are not conducive to manganese transport. Interestingly, iron levels appeared to increase in both strains after acid addition, albeit not significantly. Magnesium increased significantly at T_50_ only in the Δ*ssaACB* mutant strain. Zinc levels were slightly lower at T_50_ but this difference was not significant in either strain.

**Figure 4.**
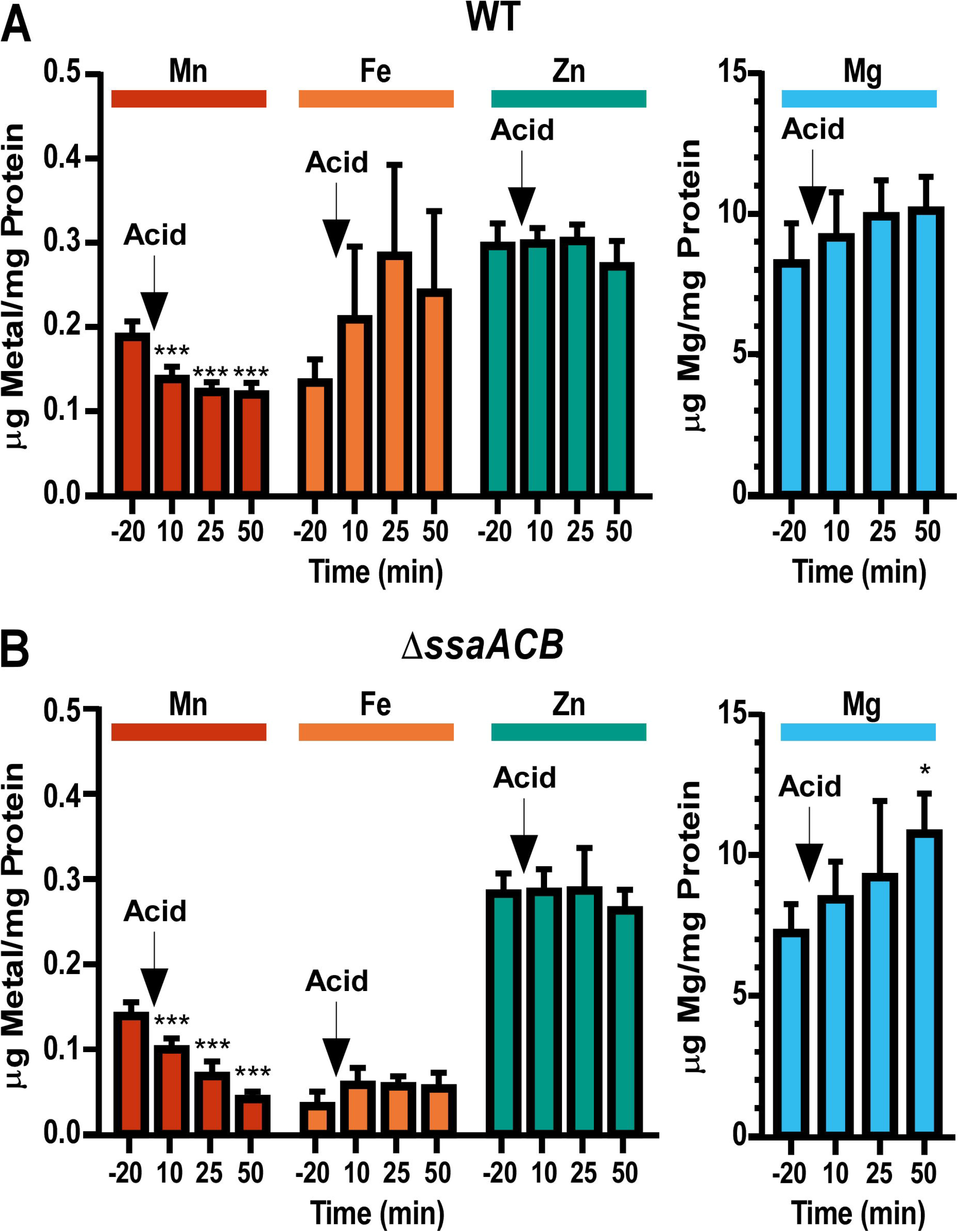
Metal content of fermentor-grown *S. sanguinis* SK36 WT and Δ*ssaACB* mutant cells before and after pH reduction. Fermentor-grown (A) WT and (B) Δ*ssaACB* cells were collected at each time point and analyzed for cellular metal content using ICP-OES. Metal concentrations were normalized to protein concentrations. Means and standard deviations of at least three replicates are displayed. Significance was determined by one-way ANOVA with Dunnett’s multiple comparisons tests with each pH 6.2 time point compared to T_−20_ (pH 7.4). \**P* ≤ 0.05, *** *P* ≤ 0.001.

To determine if reduced manganese uptake was the primary cause of the growth deficiency after pH reduction in the fermentor as it appeared to be in tube cultures, we added Mn^2+^ to a final concentration of 10 μM at T_70_ in additional fermentor runs (Figure S3). As the WT strain was not deficient in growth at pH 6.2, no restoration of growth was apparent (Figure S3A). The addition of Mn^2+^ led to a recovery in the growth rate of the Δ*ssaACB* mutant as expected (Figure S3C). When metal analysis was performed on these runs, it was observed that manganese levels 10 min after Mn^2+^ addition (T_80_) increased ~5-fold in WT (Figure S3B) but approximately doubled in the Δ*ssaACB* mutant (Figure S3D). These results suggest that Δ*ssaACB* mutant cells were still viable at the latest time point (T_50_), despite the drop in OD, since they were capable of resuming metal uptake and growth 20 min later after the addition of Mn^2+^.

### RNA-seq analysis of fermentor-grown SK36 WT and Δ*ssaACB* cells before and after pH reduction

In an attempt to determine the cause of the growth arrest of the Δ*ssaACB* mutant, we examined the transcriptome of fermentor-grown WT and Δ*ssaACB* cells taken at the same time points as described above. We examined the results of the RNA-seq analysis using three comparisons: within-strain comparisons of each pH 6.2 time point to pH 7.4 (T_−20_) for (1) WT and (2) the Δ*ssaACB* mutant (Error! Reference source not found.−S2), and (3) comparison of the WT and Δ*ssaACB* mutant strains at each time point (**Error! Reference source not found.**). The number of differentially expressed genes (DEGs; defined as *P* ≤ 0.05, |log_2_ fold change| ≥ 1) for each comparison are listed in Figure 5A. Using principal component analysis (PCA), we were able to demonstrate that each sample grouped with the others from the same time point, indicating that the results were reproducible and had minimal variability (Figure S4A-B). The 95% confidence intervals for the pH 6.2 sample time points of the WT strain overlapped with each other (Figure S4A), whereas those for Δ*ssaACB* did not (Figure S4B). The pH 7.4 samples were well separated from the pH 6.2 samples in both strains. When all samples were examined together (Figure S4C), the sample groups for both strains overlapped one another at each time point, although the WT T_50_ group overlapped more with the Δ*ssaACB* T_25_ samples than those of T_50_ (Figure S4C). These results correspond with the volcano plots (Figure S5) and heatmaps (Figure S6). Most of the expression changes occurred at the T_50_ time point for both strains, although there were fewer changes in WT than Δ*ssaACB*. When comparing the strains to each other, the T_10_ and T_25_ time points were almost identical, whereas T_−20_ and T_50_ time points had more variation (Figure 5A).

**Figure 5.**
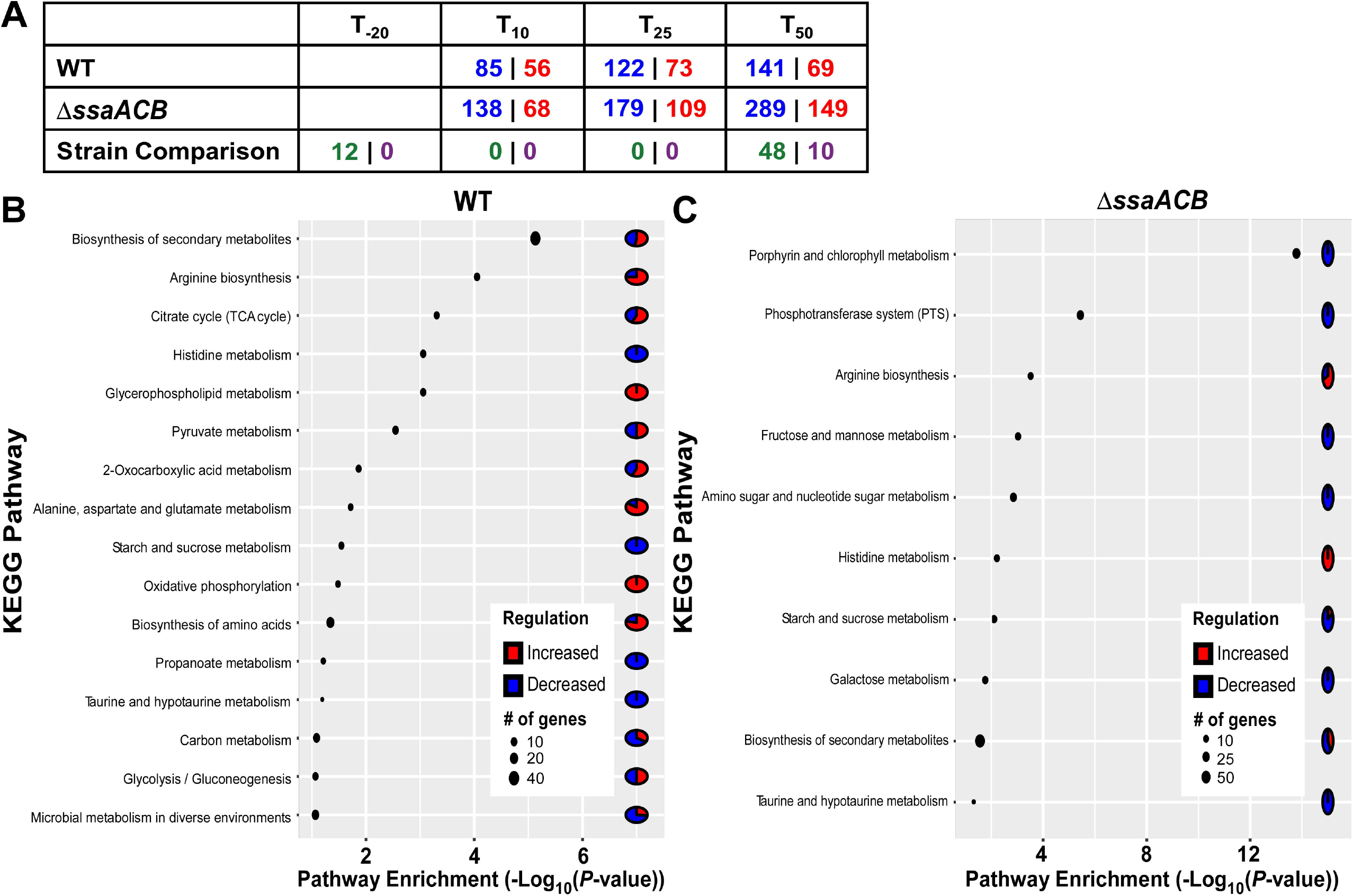
Pathway enrichment analysis of the transcriptome of fermentor-grown *S. sanguinis* SK36 WT and Δ*ssaACB* mutant cells after pH reduction. (A) Tallies of DEGs, which are defined as *P* ≤ 0.05 and |log_2_ fold change| ≥ 1. Values in blue indicate the number of genes downregulated at that time point relative to T_−20_; red values indicate those that were upregulated. Green values indicate the number of genes that were more highly expressed in WT and purple values indicate the number of genes that were more highly expressed in the Δ*ssaACB* mutant. Pathway enrichment analysis using DAVID functional classification with KEGG annotations of DEGs at T_50_ compared to T_−20_ for (B) WT and (C) Δ*ssaACB* strains.

To begin to assess the cause of the growth arrest resulting from addition of acid, we evaluated the expression of stress response genes. As expected, the expression of genes encoding an alkaline stress protein (SSA_2148), exinulcease subunit A (*uvrA*), and all subunits of the F_0_F_1_-Type ATP synthase (Kuhnert & Quivey, 2003) significantly increased in the Δ*ssaACB* mutant. Most other putative stress response genes were decreased or unchanged (**Error! Reference source not found.**). Notably, expression of the gene encoding superoxide dismutase, *sodA*, was slightly decreased. Although not unprecedented (Santi *et al.*, 2009), there is a strong connection between SodA and acid stress response in other bacteria (Kim *et al.*, 2005, Bruno-Barcena *et al.*, 2010). We presumed that the opposite response observed for *sodA* in our study was due to the reduced manganese levels in the Δ*ssaACB* mutant, because we showed in our previous study (Puccio *et al.*, 2020) that *sodA* expression is manganese dependent. Indeed, in WT cells, expression levels rose slightly. We thus considered that the unusual decrease in *sodA* expression might have contributed to the growth arrest of Δ*ssaACB* cells. We therefore tested the growth of a previously generated Δ*sodA* mutant (Crump *et al.*, 2014) in the fermentor under identical conditions, but growth was not affected (data not shown). Apart from *sodA*, these stress response genes had similar expression patterns in WT (**Error! Reference source not found.**), although changes were often of lower magnitude.

In examining the KEGG pathways assigned to the DEGs at T_50_, genes involved in phosphotransferase systems (PTS), biosynthesis of amino acids (specifically arginine and histidine), and oxidative phosphorylation were significantly affected by pH reduction in both strains. The most highly enriched pathway for the Δ*ssaACB* mutant was the “Porphyrin/Chlorophyll” pathway but upon examination, these genes encode the cobalamin biosynthetic enzymes (**Error! Reference source not found.**), which were also significantly downregulated in our recent study of manganese depletion (Puccio *et al.*, 2020). Many of the pathways highlighted by this analysis, such as carbon metabolism and amino acid metabolism, were also affected by manganese reduction in our recent transcriptomic and metabolomic studies (Puccio *et al.*, 2020, Puccio *et al.*, 2021b). We observed significant decreases in the expression of sugar transport genes (**Error! Reference source not found.**). This led us to evaluate whether acidic conditions could also lead to glucose-independent CcpA-mediated carbon catabolite repression. We examined expression of all genes found to be within the CcpA-regulon (Bai *et al.*, 2019) as well as those that we identified that have putative upstream *cre* sites (Puccio *et al.*, 2020). We found that 75 of the 382 genes with putative *cre* sites (19.6%) were significantly downregulated in the Δ*ssaACB* mutant at T_50_, whereas 30 were upregulated (7.9%) (**Error! Reference source not found.**). In WT, 47 (11.7%) were downregulated and 13 (3.4%) were upregulated. In our manganese-depletion transcriptomic study (Puccio *et al.*, 2020), we found that 19.9% of genes with putative *cre* sites were downregulated at T_50_, although the genes downregulated in the Δ*ssaACB* mutant under both conditions don’t match precisely. For example, *spxB* was significantly upregulated in reduced pH (**Error! Reference source not found.**) but significantly downregulated in reduced manganese (Puccio *et al.*, 2020). This indicates that while there may be some overlap due to the decrease in manganese levels in both studies, there are also changes that are specific to pH reduction.

As acid stress tolerance has been previously linked to amino acid biosynthesis and transport (Quivey *et al.*, 2001, Djoko *et al.*, 2017, Guan & Liu, 2020, Senouci-Rezkallah *et al.*, 2011), we evaluated the expression of genes annotated with these functions (**Error! Reference source not found.**). Many of these genes were significantly affected by acid addition, although more were affected in the Δ*ssaACB* mutant. Of note, aconitate hydratase, citrate synthase, and isocitrate dehydrogenase were significantly upregulated in the Δ*ssaACB* mutant but significantly downregulated in the WT strain at T_50_ (**Error! Reference source not found.**).

### Biofilm competition of the *S. sanguinis* SK36 WT and Δ*ssaACB* strains against *S. mutans in vitro*

After examining the effect of reduced manganese on gene expression and loss of manganese transporters on growth and metal content, we decided to assess whether the lack of the primary manganese transporter would have relevance to growth conditions similar to those found in the oral cavity. We therefore compared the growth of a spectinomycin (Spc) resistant SK36 WT strain, JFP56, to the Δ*ssaACB* mutant in competition with *S. mutans* (Figure 6). Given that *S. mutans* often outcompetes WT *S. sanguinis* if given the opportunity to colonize first, we inoculated wells of biofilm media (BM) + 1% sucrose with *S. sanguinis* to stimulate biofilm formation and allowed 24 h of aerobic growth. We then refreshed the wells with Tryptone Yeast Extract + 1% sucrose (TY+S) and inoculated with an *S. mutans* strain containing a plasmid-encoded erythromycin (Erm) resistance gene. After another 24 h incubation, we then measured the pH of the media and plated the biofilm cultures. *S. mutans* growth was unaffected by the presence of *S. sanguinis* (Figure 6A), whereas both *S. sanguinis* strains were recovered in significantly lower numbers in the presence of *S. mutans* (Figure 6B). In fact, only one of six replicates of the Δ*ssaACB* mutant produced a single colony after incubation in the presence of *S. mutans.* The pH of all culture supernatants after 24 h growth in TY+S was significantly lower than media alone, with the *S. mutans* supernatant averaging pH 4.08 and the *S. sanguinis* WT and Δ*ssaACB* supernatants averaging pH 4.50 and 4.58, respectively (Figure 6C).

**Figure 6.**
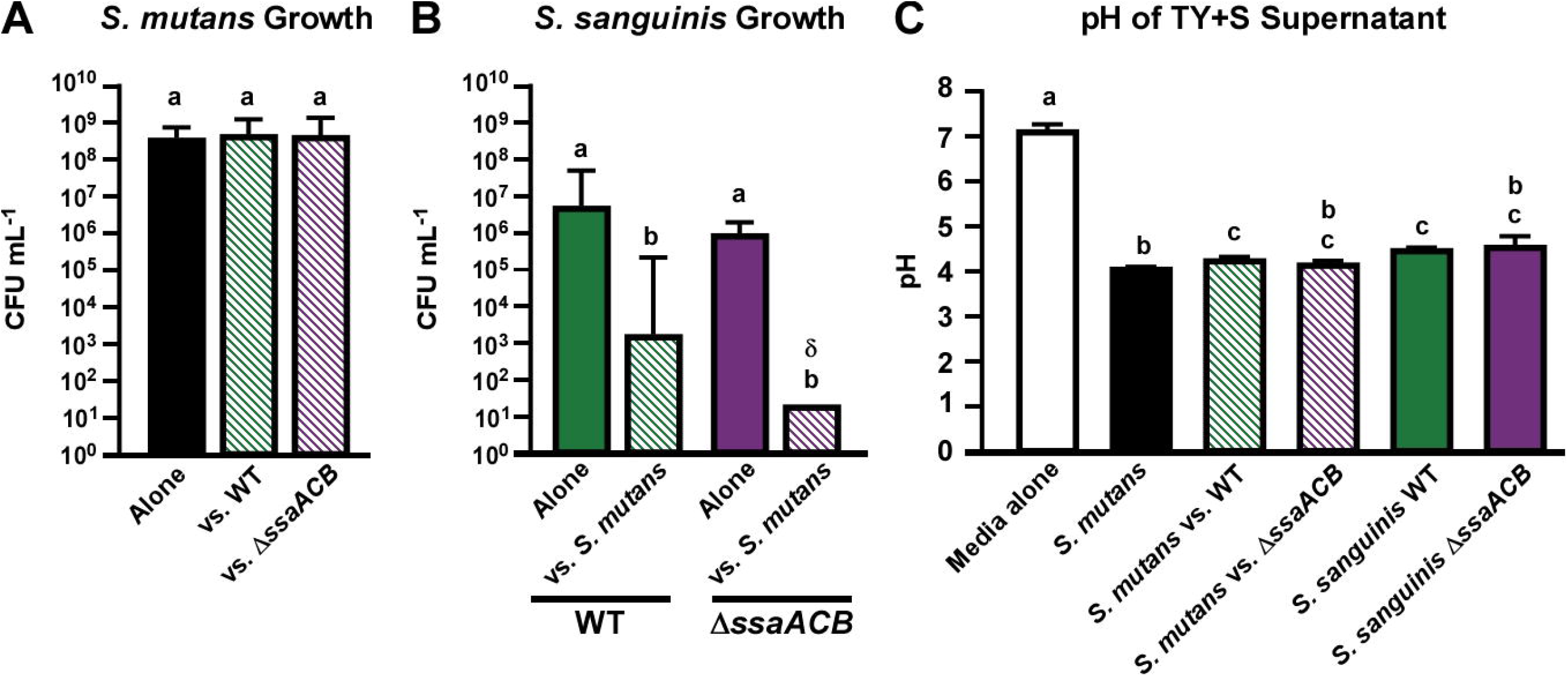
Competition of *S. sanguinis* SK36 WT and Δ*ssaACB* strains against *S. mutans in vitro*. *S. sanguinis* cultures were grown aerobically for 24 h in BM+S. Media was swapped for TY+S and *S. mutans* was inoculated. Cultures were incubated an additional 24 h before media was removed and cells scraped from wells were resuspended in PBS for plating on BHI agar plates with antibiotics for selection. Means and standard deviations of *S. mutans* (A) and *S. sanguinis* (B) CFU mL^−1^ from three biological replicates with two technical replicates each are shown. The δ indicates that 5 of 6 replicates fell below the limit of detection. (C) Means and standard deviation of media pH measurements for each culture are shown. Statistical analysis was performed using one-way ANOVA with a Tukey multiple comparisons post-test. Bars with the same letter within a chart are not significantly different (*P* > 0.05).

### Contribution of SsaACB to oral colonization and competition in a murine model

Given the results of the *in vitro* competition assay, we used an *in vivo* model to compare colonization of the oral cavity and dental biofilms by the SK36 WT and Δ*ssaACB* mutant strains, and once colonization was established, we then compared the ability of each strain to compete against *S. mutans* under cariogenic conditions. We employed the mouse model described recently by Culp *et al.* (2020) with mice fed a diet containing 37.5% sucrose. After antibiotic suppression of the oral microbiota, each of two groups of mice received five daily oral inoculations with one of the two *S. sanguinis* test strains, followed two weeks later by challenge with *S. mutans*. Oral swabs were taken at weekly intervals, thus providing a measure of colonization within saliva and the oral mucosal pellicle, with subsequent sonication of mandibular molars to assess bacteria recovered from dental biofilms (see timeline of events, Figure 7A). Recoveries of bacteria released from swabs and molar sonicates were determined by strain-specific qPCR assays of genomic DNA, plus estimates of total recovered bacteria were determined by qPCR assay of the ubiquitous single copy gene, *rpsL* (30S ribosomal protein S12). Subtraction of strain-specific recoveries from total recovered bacteria allowed estimations of the population of recovered murine oral commensal bacteria. We observed that both *S. sanguinis* strains colonized well in competition with native mouse commensals (Figure 7B). Upon introduction of *S. mutans*, the abundance of the two test strains and the mouse commensals declined and then recovered by the following week. When levels of the two test strains were compared for each time point, we found that the Δ*ssaACB* mutant was recovered from the molars at slightly, yet significantly lower levels than WT 14 days after the introduction of *S. mutans* (experimental day 28). The mouse commensal species present in each animal group were also compared to one another at each time point and were only found to be significantly different on the molars of mice inoculated with the Δ*ssaACB* mutant. *S. mutans* levels were slightly but not significantly higher in competition with SK36 WT at day 27 (swab 5) than in competition with the Δ*ssaACB* mutant on the same day. These results suggest that SsaACB had no impact on oral colonization and was of only minor importance for dental colonization and competition, even in a strain that did not possess the *mntH* gene. This is in stark contrast to the results obtained from our *in vivo* endocarditis model showing that the SsaACB transporter is essential for virulence, even in strains possessing TmpA and MntH transporters (Puccio *et al.*, 2021a).

**Figure 7.**
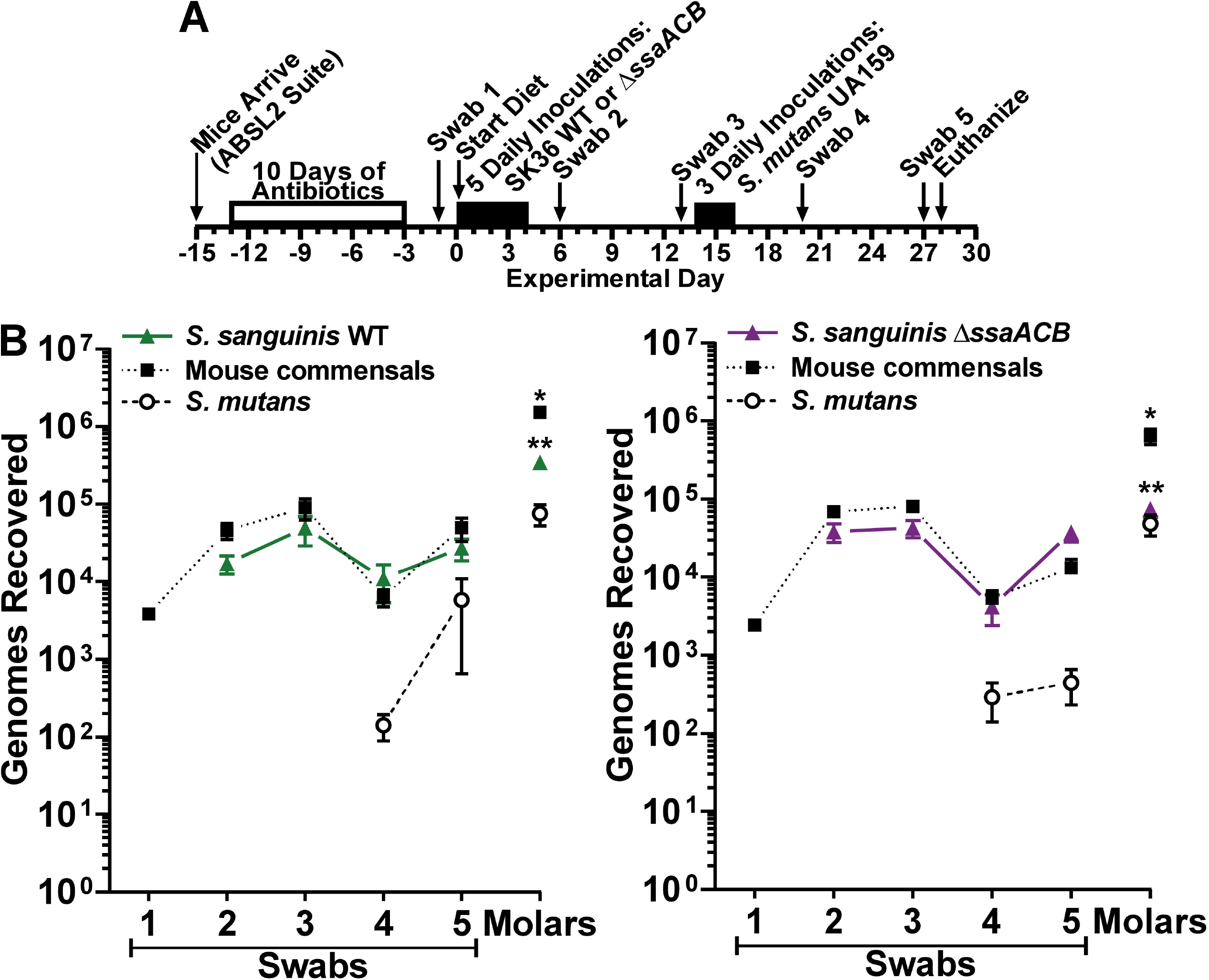
Comparison of oral colonization by *S. sanguinis* SK36 WT and Δ*ssaACB* strains, and in subsequent competition with *S. mutans* UA159 *in vivo*. (A) Timeline of key events in the experiment. (B) Colonization results for each indicated *S. sanguinis* strain (closed triangles, solid lines), *S. mutans* UA159 (open circles, dashed lines) and mouse oral commensals (closed squares, dotted lines) from oral swabs 1-5 taken at the times indicated in A and from sonicates of mandibular molars. *S. sanguinis* SK36 WT is depicted in green (left) and the Δ*ssaACB* mutant is depicted in purple (right). Means and standard error (n = 12 mice per cohort) of recovered genomes estimated by qPCR are shown. Significant differences within each bacterial species/group between the two cohorts of mice were calculated using one-way ANOVA with Bonferroni’s multiple comparisons tests for each sample time point (* *P* < 0.05, ** *P* < 0.01). Mice were fed a high-sucrose powdered diet with sterile drinking water.

## Discussion

While a relationship between manganese and acid tolerance in streptococci has been suggested previously (Shabayek & Spellerberg, 2017, Kajfasz *et al.*, 2020), it has yet to be fully characterized. Beighton (1982) noted that manganese induced *S. mutans* to form caries and influenced carbohydrate metabolism. In *Streptococcus pneumoniae* (Martin-Galiano *et al.*, 2005) and *S. agalactiae* (Santi *et al.*, 2009), the orthologs of SsaACB were upregulated in response to acid stress. In *S. mutans*, expression of the manganese-dependent regulator SloR was previously linked to ATR (Dunning *et al.*, 2008). Recently, it was appreciated that loss of MntH in *S. mutans* (Kajfasz *et al.*, 2020) and *S. agalactiae* (Shabayek *et al.*, 2016) led to a reduction in acid tolerance. Here we report that loss of the high-affinity manganese ABC transporter in *S. sanguinis* strain SK36, SsaACB, reduces acid tolerance but does not greatly influence oral colonization or competition with *S. mutans*. We also characterize the transcriptional changes that accompany the growth arrest of the Δ*ssaACB* mutant observed in reduced pH conditions *in vitro* and in comparison with WT.

Initially, we were unsure whether the growth arrest we observed in the Δ*ssaACB* mutant was directly related to manganese, but the reduction in intracellular manganese levels in both WT and Δ*ssaACB* strains in tube cultures (Figure 3) and fermentor cultures (Figure 4) indicates that reduced pH affects the ability of manganese to enter the cell. No other biologically relevant metal significantly decreased in cells at pH 6.2 (Figure S1), further highlighting the relationship between manganese and reduced pH. Mean iron levels increased in both strains in the fermentor after pH reduction, albeit not significantly (Figure 4). These correspond to increases in the expression of genes encoding putative iron transporters (Tables S1-S2), such as those encoded by SSA_RS07705-SSA_RS07715. Similarly, the significant increase in magnesium levels in the Δ*ssaACB* mutant may be due to an increase in expression of SSA_RS03540 (Table S2), which encodes a putative magnesium-transport protein, CorA (Kehres *et al.*, 1998). Moreover, the addition of as little as 1 μM Mn^2+^ restored the growth of the Δ*ssaACB* mutant to WT-like levels, suggesting that this is likely the main reason for the growth arrest observed at pH 6.2. Furthermore, many transcriptomic responses to pH reduction were similar to those of manganese reduction by EDTA (Puccio *et al.*, 2021a).

We hypothesized that either the function of a secondary manganese transporter, TmpA, is negatively affected by reduced pH or that bioavailability of manganese is affected. The ortholog of TmpA in *E. coli*, ZupT, functions best near neutral pH (Taudte & Grass, 2010). Several lines of evidence in this study suggest that at pH 6.2, TmpA loses the ability to transport manganese efficiently. We found that manganese levels in a Δ*ssaACB* Δ*tmpA* mutant strain were indistinguishable from those in the Δ*ssaACB* parent in BHI at pH 6.2 ± 10 μM Mn^2+^ (Figure 3B), which was not true at pH 7.3 (Figure 3A). Moreover, manganese levels of both strains at pH 6.2 were very similar to levels in the Δ*ssaACB* Δ*tmpA* mutant at pH 7.3, suggesting that growth at pH 6.2 has an effect equivalent to deletion of the *tmpA* gene. Additionally, growth of the SK36 Δ*ssaACB* and Δ*ssaACB* Δ*tmpA* mutants were indistinguishable at pH 6.2 and reached a final density that was similar to that of the Δ*ssaACB* Δ*tmpA* mutant at pH 7.3, again suggesting that growth at pH 6.2 has the same effect as deletion of the *tmpA* gene. Finally, when strains that possess a NRAMP protein (MntH), such as VMC66, are deleted for *ssaACB*, they do not experience a decrease in manganese levels or growth at pH 6.2, in agreement with expectations if one assumes the activity of TmpA is greatly diminished at this pH and that of MntH is not. Taken together, these results strongly support the hypothesis that reductions in growth and manganese levels of SK36 and its Δ*ssaACB* mutant at pH 6.2 are due to the reduced function of TmpA (Figure 8), although studies with cell-free liposomes will be required for confirmation.

**Figure 8.**
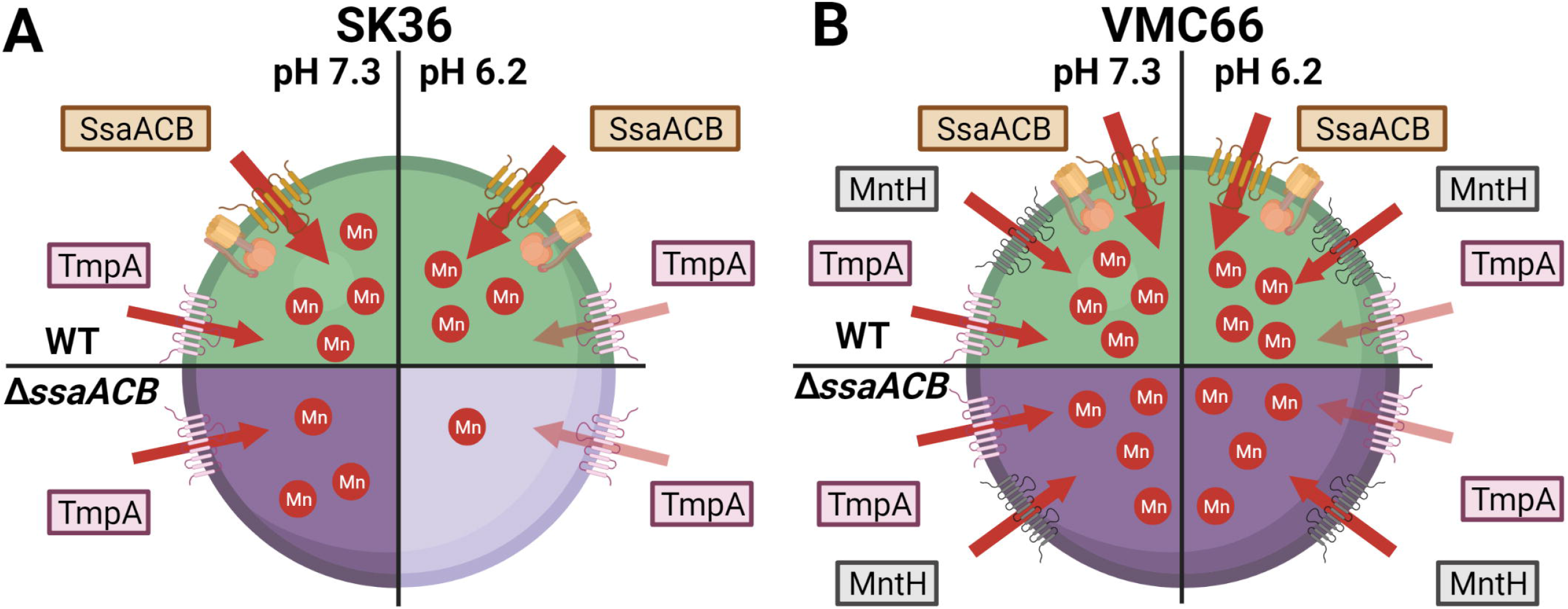
Summary of the role of manganese transporters in the growth of *S. sanguinis* in reduced-pH conditions. Diagrams of *S. sanguinis* WT (A) SK36 and (B) VMC66 strains at pH 7.3 (left) and pH 6.2 (right) with their known manganese transporters depicted. In both WT cells (top), all transporters are present so although TmpA function is reduced, manganese is acquired in sufficient levels to support growth. In SK36 Δ*ssaACB* mutant cells (bottom), TmpA is capable of transporting enough manganese at pH 7.4 to support the fast growth rate of cells in the fermentor; however, at pH 6.2, TmpA-mediated manganese transport is insufficient to maintain growth in the fermentor or in tube cultures. In VMC66 Δ*ssaACB* mutant cells (bottom), MntH is able to compensate for the reduced function of TmpA at pH 6.2, resulting in sufficient manganese levels to support growth. Either TmpA or MntH can provide sufficient manganese to support growth at pH 7.3 in the absence of the other two transporters; this is also true at pH 6.2 for MntH but not for TmpA.

Due to the short time frame of our transcriptional analysis (50 min), we believe that the changes observed are an acid shock response, as opposed to acid adaptation (Quivey *et al.*, 2001). However, acid shock responses have typically employed a pH of 3.0 to 4.4 (Martin-Galiano *et al.*, 2005, Hamilton & Svensäter, 1998) so given that the growth rate of WT is unchanged and that we used a modest reduction in pH (6.2), it is likely a relatively mild response. When comparing the reduced-pH RNA-seq results in each strain directly to the other, few DEGs were observed and those that were occurred only at the T_−20_ and T_50_ time points (Figure 5A). Despite these results, we observed a drastic difference in the fermentor growth rate between WT and Δ*ssaACB* after acid addition (Figure S2). Thus, we hypothesize that either subtle changes between strains have a large combined effect or post-transcriptional changes not captured by transcriptomics may be influencing the growth. It is also possible that both explanations are correct. One reason for the changes being small could be that WT and Δ*ssaACB* are both experiencing two, typically separate stresses—acid stress and manganese depletion—although SK36 is experiencing both to a lesser degree. Examination of the manganese levels of WT cells in unsupplemented batch cultures at pH 7.3 vs. 6.2 (Figure 3A vs. 3B) and fermentor cultures before and after acid addition (Figure 4A) suggests that even with SsaACB intact, WT cells take up less manganese at pH 6.2 than 7.3. This is also supported by the increase in expression of *ssaB*, *ssaC*, and *ssaA* (SSA_RS01450-01460) after acid addition in the fermentor (Table S1). However, it is also clear that both strains exhibit expression changes expected in response to reduced pH, as discussed previously. It would not be surprising if the effects of the two stresses were additive or synergistic with regard to growth inhibition. Thus, expression changes would be spread over two stress responses rather than one, further diminishing the importance or magnitude of any one expression difference.

We recently reported that manganese depletion led to induction of carbon catabolite repression in *S. sanguinis* (Puccio *et al.*, 2020, Puccio *et al.*, 2021b). Given that both addition of acid and EDTA (Puccio *et al.*, 2021b) led to reduced manganese levels in the fermentor (Figure 4), we hypothesize that the impact on glycolysis is similar to what we observed in our previous study. Additionally, expression of genes encoding glycolytic enzymes responded similarly in both studies (**Error! Reference source not found.**), indicating that the regulatory mechanisms controlling the expression of these genes may have a manganese-related aspect. Thus we hypothesize that there is likely an accumulation of FBP occurring in cells growing at reduced pH due to reduced activity of Fba, Fbp, or both enzymes (Puccio *et al.*, 2021b). This may be the cause behind the changes in expression we observed in the CcpA regulon (**Error! Reference source not found.**). As noted by Radin *et al.* (2016), glycolytic enzymes may have a higher demand for manganese than other enzymes and thus in manganese-deplete conditions, the cells may shift to amino acid metabolism as a source of energy. This could explain why we observed significant increases in expression of many amino acid biosynthetic and transport genes (**Error! Reference source not found.**). However, the impact of PTS, CcpA, and central carbon metabolism on gene expression are complex, and it is likely that multiple factors impact the expression of CCR-responsive genes.

Acid is an important component of the competition between *S. sanguinis* and *S. mutans in vitro* (Kreth *et al.*, 2005). While not significant, the difference in the mean levels of SK36 WT and Δ*ssaACB* mutant growth in direct competition with *S. mutans* in *in vitro* biofilms was appreciable, especially given that the Δ*ssaACB* mutant was only recovered in one out of six replicates. Our *in vivo* study yielded no significant differences in recoveries from oral swabs, a measure of colonization of oral mucosal epithelia (Culp *et al.*, 2020), and only slight, yet significant differences between the SK36 WT and Δ*ssaACB* mutant strains in colonization of dental biofilms when in competition with *S. mutans*. It was recently shown using the same mouse model that clinical isolates of streptococci from dental plaque of caries-free individuals (*i.e.*, *Streptococcus mitis*, *Streptococcus gordonii*, *Streptococcus* A12), which *in vitro* were highly competitive against *S. mutans*, poorly colonized oral mucosal and dental biofilms, while two strains of *S. sanguinis* suppressed initial mucosal colonization by *S. mutans* and persisted in both biofilms, even as *S. mutans* recovered to levels observed in a control group of mice challenged with *S. mutans* alone (Culp *et al.*, 2020). Moreover, both strains of *S. sanguinis* promoted dental colonization of mouse commensals to levels equivalent or higher than *S. mutans*, similar to the SK36 WT and Δ*ssaACB* mutant strains in the current study. Additionally, oral colonization of SK36 WT and its Δ*ssaACB* mutant quickly recovered after being initially suppressed when challenged with *S. mutans*, suggesting that persistent colonization of mucosal biofilms may serve as a reservoir for colonization of dental biofilms. In clinical studies, *S. sanguinis* was shown to precede and delay colonization by *S. mutans* of newly acquired teeth in children (Caufield *et al.*, 2000) and to persist in cavitated lesions in which *S. mutans* accounted for up to 55% of the microbiota (Gross *et al.*, 2012). Collective evidence therefore indicates *S. sanguinis* is well adapted to the oral cavity and dental biofilms in which it copes with innate immune factors in saliva, interacts with other oral commensals either competitively or cooperatively, and importantly can coexist with *S. mutans* under cariogenic conditions. Although the SsaACB transport system appears to be universally conserved in *S. sanguinis* strains, its importance was not demonstrable under *in vivo* conditions.

We propose three possible explanations for the *in vivo* results. One is that SsaACB wasn’t necessary because the oral environment contained sufficient manganese or had a pH close enough to neutral to allow comparable growth of the WT and Δ*ssaACB* strains using only the TmpA transporter over the course of this experiment. Secondly, as described above from the work of Radin *et al.* (2016), a shift to amino acid metabolism as a source of energy may have lessened the demand for manganese under conditions of manganese depletion. It has been shown that interactions of oral commensal species and *S. mutans* with surface-associated salivary constituents induces a more protease-active phenotype capable of more effectively degrading salivary proteins and mucin glycoproteins (Kindblom *et al.*, 2012, Wickström *et al.*, 2013), thus potentially providing a reliable source of amino acids for uptake and metabolism. Another is that perhaps there is some aspect of the oral cavity environment that reduces the activity or expression of SsaACB. In this case, SsaACB would not be capable of making a strong contribution, so the WT strain would compete only slightly better than the mutant. While this explanation seems inherently less plausible, we cannot rule it out at present. If true, it could account for the existence of TmpA in *S. sanguinis*, which is otherwise inexplicable. Moreover, although we have not yet tested the effect of MntH in an oral model, the fact that it seems to be most active under exactly the conditions in which TmpA is least active (Figure 8) provides a rationale for its presence in some strains.

In conclusion, these results indicate that SsaACB made no more than a minor contribution to competitiveness in this model, despite the fact that the only other transporter, TmpA, has now been found to be inactive at reduced pH. These results may guide future studies. For example, there has been much interest concerning the development of probiotic bacteria for promotion of oral health. Given the results of the studies employing the endocarditis model in the accompanying paper (Puccio *et al.*, 2021a) in combination with the murine studies reported here, and the work with the Δ*mntH* mutant of VMC66, one course of action would be to begin with a promising *mntH-*containing *S. sanguinis* strain or candidate probiotic species such as *Streptococcus* sp. A12 (Lee *et al.*, 2019), then delete *ssaACB*. This mutant could be tested in a similar manner to the experiments described here and in the accompanying paper to ensure the same behavior of the mutants. Additionally, these data suggest that an SsaB inhibitor may be effective at preventing IE while leaving the oral flora intact. Therefore, an IE patient with a healthy oral microbiota could take a small-molecule SsaB inhibitor to prevent IE with less concern of shifting the oral microbiota in favor of caries. Furthermore, IE patients with high risk for dental caries could take a probiotic in addition to the SsaB inhibitor without additional risk of IE. In conclusion, these results and those of the accompanying study (Puccio *et al.*, 2021a) enhance the understanding of the role of manganese in *S. sanguinis* and highlight this essential nutrient as an important factor for consideration in the development of streptococcal therapeutics.

### Experimental Procedures

#### Bacterial strains

Strains and plasmids used in this study are listed in Table 1. All strains were grown in overnight cultures from single-use aliquots of cryopreserved cells, diluted 1000-fold in BHI media (Beckinson Dickinson). Mutant strains were incubated with the appropriate antibiotics: kanamycin (Kan; Sigma-Aldrich) 500 ug mL^−1^; erythromycin (Erm; Sigma-Aldrich); 10 μg mL^−1^; spectinomycin (Spc; Sigma-Aldrich) 200 μg mL^−1^; chloramphenicol (Cm; Fisher) 5 μg mL^−1^. The pre-cultures of the SK36 Δ*ssaACB* Δ*tmpA* and VMC66 Δ*ssaACB* Δ*mntH* mutants required exogenous Mn^2+^ (10 μM) for growth. The cultures were then incubated at 37°C for 16-20 h with the atmospheric condition set to 1% O_2_ (9.5% H_2_, 9.5% CO_2_, and 80% N_2_) or 6% O_2_ (7% H_2_, 7% CO_2_, and 80% N_2_) using a programmable Anoxomat™ Mark II jar-filling system (AIG, Inc).

**Table 1.** Strains and plasmids used in this study.

| Strain | Description | Source or Reference |
| --- | --- | --- |
| SK36 | Human oral plaque isolate | Mogens Killan (Aarhus University); Xu *et al.* (2007) |
| JFP56 | ΔSSA_0169::*aad9*, derived from SK36 | Turner *et al.* (2009) |
| JFP169<sup>†</sup> | ΔssaACB::aphA-3, derived from SK36 | Baker *et al.* (2019) |
| JFP173<sup>‡</sup> | ΔssaACB::tetM, derived from SK36 | Baker *et al.* (2019) |
| JFP226 | ΔtmpA::aphA-3, derived from SK36 | Puccio *et al.* (2021a) |
| JFP227<sup>§</sup> | ΔssaACB::tetM ΔtmpA::aphA-3, unintentional mutation: SSA_1414-W139*; derived from SK36 | Puccio *et al.* (2021a) |
| JFP377 | ΔssaACB::tetM ΔtmpA::aphA-3, clean version, derived from SK36 | Puccio *et al.* (2021a) |
| VMC66 | Human endocarditis isolate | Kitten *et al.* (2012) |
| JFP313 | ΔssaACB::aphA-3, derived from VMC66 | Puccio *et al.* (2021a) |
| JFP317 | ΔtmpA::pSerm, derived from VMC66 | Puccio *et al.* (2021a) |
| JFP320 | ΔssaACB::aphA-3 ΔtmpA::pSerm, derived from VMC66 | Puccio *et al.* (2021a) |
| JFP367 | ΔmntH::tetM, derived from VMC66 | Puccio *et al.* (2021a) |
| JFP370 | ΔssaACB::cat ΔmntH::tetM, derived from VMC66 | Puccio *et al.* (2021a) |
| UA159 | Wild type *S. mutans* | ATCC 700610 |
| Plasmid | Description | References |
| pVMTeal | Plasmid encoding Erm<sup>R</sup> for use in *S. mutans* | Vickerman *et al.* (2015)<br>Zhu *et al.* (2017) |

<sup>†</sup> Designated $\Delta$ ssaACB throughout the manuscript.
<sup>‡</sup> Used only for growth studies and metal analysis in Figures 2, 3, and S1.
<sup>§</sup> Used only for metal analysis in Figure 3A.

### Tube growth studies

The pH of BHI was modified by addition of 6 N HCl prior to autoclaving. Each tube was preincubated at 37°C in 1% O_2_ in an Anoxomat jar overnight, then inoculated with a 10^−6^-fold dilution of the overnight pre-culture. The inoculated tubes were returned to incubate at 37°C in 1% O_2_. In some growth studies, MnSO_4_ (Alfa Aesar; Puratronic™, >99.999% pure) was added to culture tubes immediately prior to inoculation. To determine CFUs, samples were sonicated for 90 s using an ultrasonic homogenizer (Biologics, Inc) to disrupt chains. Cultures were diluted in PBS and plated on BHI agar (Beckinson Dickinson) plates using an Eddy Jet 2 spiral plater (Neutec Group, Inc.). Plates were incubated for 24 h at 37°C under anaerobic conditions with a palladium catalyst included in the jars.

### Metal analysis

For metal analysis of cells growing batch cultures, 2 tubes of 38 mL of BHI per strain and condition were pre-incubated at 37°C overnight. For some samples, Mn^2+^ was added to 10 μM immediately prior to inoculation. Pre-cultures (3 mL) were added to each tube. For pH 7.3 samples, pre-cultures were added directly. For pH 6.2 samples, pre-cultures were centrifuged and resuspended in pH 6.2 BHI prior to inoculation. Cultures were grown to mid-logarithmic phase and centrifuged at 3,740 × *g* for 10 min at 4°C. All strains grew under the conditions of this assay; however, some of the mutants grew slower than the WT strains. For fermentor samples, aliquots of 40 mL were collected at each time point and then centrifuged as described above.

The collected cell pellet was then washed twice with cold cPBS (PBS treated with Chelex-100 resin (Bio-Rad) for 2 h, then filter sterilized and supplemented with EDTA to 1 mM). The pellet was then divided for subsequent acid digestion or protein concentration determination. Trace metal grade (TMG) nitric acid (15%) (Fisher Chemical) was added to one portion of the pellet. The pellet was digested using an Anton Paar microwave digestion system using a modified Organic B protocol: 120°C for 10 min, 180°C for 20 min, with the maximum temperature set to 180°C. The digested samples were then diluted 3-fold with Chelex-treated dH_2_O. Metal concentrations were determined using an Agilent 5110 inductively coupled plasma-optical emission spectrometer (ICP-OES). Concentrations were determined by comparison with a standard curve created with a 10 μg ml^−1^ multi-element standard (CMS-5; Inorganic Ventures) diluted in 5% TMG nitric acid. Pb (Inorganic Ventures) was used as an internal standard (100 μg ml^−1^). The other portion of the pellet was resuspended in PBS and mechanically lysed using a FastPrep-24 instrument with Lysing Matrix B tubes (MP Biomedicals) as described previously (Rhodes *et al.*, 2014). Insoluble material was removed by centrifugation. Protein concentrations were determined using a bicinchoninic acid (BCA) Protein Assay Kit (Pierce) as recommended by the manufacturer, with bovine serum albumin as the standard.

### Fermentor growth conditions

The fermentor conditions were similar to Puccio and Kitten (2020) with minor modifications. A BIOSTAT® B bioreactor (Sartorius Stedim) with a 1.5-L capacity UniVessel^®^ glass vessel was used for growth of 800-mL cultures at 37°C. Cultures were stirred at 250 rpm and pH was maintained by the automated addition of 2 N KOH and 2 N HCl (Fisher Chemical). A 40-mL overnight pre-culture of *S. sanguinis* was grown as described above and centrifuged for 10 minutes at 3,740 *x g* in an Allegra X-142 centrifuge at 4°C. The supernatant was discarded and the cells were resuspended in BHI prior to inoculation. The air flow was kept at 0.03 lpm for the entire experiment. Once the cells reached their peak OD in static culture, the input flow of BHI was set to 17% (~700 mL h^−1^), and the output flow of waste was set to 34% for the remainder of the experiment. Cells were allowed to acclimate to this media flow rate for 1 h. The T_−20_ sample was aseptically removed for total RNA isolation or metal analysis. The fermentor culture in the vessel was adjusted to pH 6.2 using an in-dwelling probe at T_0_. Samples were taken for each post-treatment time point (T_10_, T_25_, T_50_). In some experiments, MnSO_4_ (Puratronic™; Alfa Aesar) was added to the carboy (T_66_) and vessel (T_70_) at a final concentration of 10 μM and samples were taken for metal analysis at T_80_.

### RNA isolation

Fermentor samples (2 mL) were added to 4 mL RNAprotect Bacteria Reagent (Qiagen) and immediately vortexed for 10 s. The samples were then incubated at room temperature for 5-90 min and centrifuged for 10 min at 3,740 × *g* at 4°C. The supernatant was discarded and the samples stored at −80°C. RNA isolation and on-column DNase treatment were completed using the RNeasy Mini Kit and RNase-Free DNase Kit, respectively (Qiagen). RNA was eluted in 50 μL RNase-Free water (Qiagen). A second DNase treatment was then performed on the samples (Invitrogen).

### RNA sequencing

Total RNA quantity and integrity were determined using a Bioanalyzer (Agilent). All samples passed quality control assessment with RNA Integrity Numbers (RIN) above 8. Two sequential rounds of ribosomal reduction were then performed on all samples using RiboMinus™ Transcriptome Isolation Kit (ThermoFisher). The resulting depleted RNA was assessed using Bioanalyzer (Agilent) to confirm efficient rRNA removal. Stranded RNA-seq library construction was then performed on the rRNA-depleted RNA using the Kapa RNA HyperPlus kit for Illumina (Roche) following manufacturer’s specifications for library construction and multiplexing. Final Illumina libraries were assessed for quality using an Agilent Bioanalyzer DNA High Sensitivity Assay and qPCR quantification was performed using Kapa Library Quantification kit for Illumina (Roche). Individual libraries were pooled equimolarly and the final pool was sequenced on an Illumina MiSeq with 2 × 75-bp paired-end reads. Demultiplexing was performed on the Illumina MiSeq’s on-board computer. The Virginia Commonwealth University Genomics Core Facility completed all RNA-seq library preparation and sequencing.

### RNA-seq analysis pipeline

Using Geneious Prime 2021.1.1 (https://www.geneious.com), sequence reads were trimmed using the BBDuk Trimmer prior to mapping to either the SK36 genome or a modified version, in which the *ssaACB* operon was replaced with the *aphA-3* sequence. Original and new locus tags from the Genbank® annotations are included (Benson *et al.*, 2013). PATRIC annotations (https://patricbrc.org/) (Wattam *et al.*, 2017) are also included. Reads for each post-treatment sample were compared to the corresponding pre-treatment (T_−20_) sample using DESeq2 (Love *et al.*, 2014) in Geneious to determine log_2_ fold changes and adjusted *P*-values. The same method was used to compare reads from WT to those from the Δ*ssaACB* mutant at each sample time point. Principal component analysis was completed using R (version 4.0.5) and RStudio (version 13.959) with Bioconductor (Bioconductor.com) package pcaExplorer version 2.160 (Marini & Binder, 2019). Volcano plots were generated using R and RStudio with Bioconductor package EnhancedVolcano version 1.8.0 (Blighe *et al.*, 2018). All DEGs were input into the DAVID database (https://david.ncifcrf.gov/summary.jsp) (Dennis *et al.*, 2003). Since the new locus tags are not accepted in KEGG, only genes with original “SSA_” locus tags were included in the analysis. The KEGG_pathway option was chosen for functional annotation clustering. The *P*-value shows the significance of pathway enrichment. DAVID pathway figures were generated using an R script (https://github.com/DrBinZhu/DAVID_FIG).

### Biofilm competition assays

*S. sanguinis* pre-cultures (described above) were diluted 100-fold into 2 mL pre-warmed BM (Loo *et al.*, 2000) + 1% sucrose in 12 well plates. The cultures were grown for 24 h at 37°C aerobically (~21% O_2_). The media supernatant was carefully removed and discarded. Warm TY (3% tryptone, 0.1% yeast extract, 0.5% KOH, 1 mM H_3_PO_4_) (Fozo & Quivey, 2004) + 1% sucrose was added to 2 mL and *S. mutans* + pVMTeal pre-cultures were diluted 100-fold into the wells. After 24 h incubation at 37°C, TY+S supernatant was carefully removed and the pH was measured. PBS was added to 1 mL and biofilms were scraped from each well. Each biofilm culture was sonicated for 90 s using an ultrasonic homogenizer. Cultures were diluted in PBS and plated on BHI plates with antibiotics using a spiral plater. Plates were incubated at 37°C for 24 h at 0% O_2_.

### Mouse model of oral colonization

The mouse model was described in detail recently by Culp *et al.* (2020). Briefly, all procedures with solutions and samples were performed under BSL2 conditions and mice were kept under ABSL2 conditions. Inbred 3-week-old female SPF BALB/cJ mice (The Jackson Laboratory, Bar Harbor, ME) were placed in pairs in sterile cages. Two days later, mice were given drinking water containing 0.8 mg/ml sulphamethoxazole/0.16 mg/ml trimethoprim for a total of 10 days to suppress indigenous oral bacteria, followed by a 3-day washout period with sterile drinking water. On the following day (designated experimental day 0) mice were placed on custom diet TD.160810 (Teklad, Madison, WI) containing 37.5% sucrose and without fluoride, and inoculated daily over five days with 50 μl of 1.5% (wt/vol) carboxymethylcellulose in saliva buffer (50 mM KCl, 1.0 mM KPO_4_, 0.35 mM K_2_HPO_4_, 1.0 mM CaCl_2_ 2H_2_O, 0.1 mM MgCl_2_ 6H_2_O, pH 6.5) containing approximately 1×10^9^ cells of the indicated strain grown to an OD_600_ between 0.55 to 0.70 in BHI. Two weeks later, mice received three daily inoculations with approximately 1×10^9^ cells of *S. mutans* UA159. Mice were euthanized by CO_2_ asphyxiation followed by cervical dislocation. The protocol was approved by the Institutional Animal Care and Use Committee at University of Florida (IACUC protocol #201810470). Oral swabs were taken using HydraFlock® 6” Sterile Micro Ultrafine Flock swabs (Puritan Medical Products, Guilford, ME). Swab tips were vortexed (3 times for 5 sec) in 1 ml sterile PBS, the tips removed and 200 μl added of ice-cold PBS containing approximately 5 × 10^8^ depurinated cells of laboratory strain *S. mitis* UF2. Tubes were then vortexed 5 sec and centrifuged (10,000 × *g*, 10 min at 4 °C) to pellet recovered cells. Cell pellets were then processed for DNA isolation using the DNeasy UltraClean Microbial kit (Qiagen Inc., Germantown, MD) as per manufacturer’s instructions. Depurinated cells were devoid of detectable DNA by qPCR and allowed for quantitative recoveries of DNA from test strains and mouse commensals. To measure dental colonization, the left and right halves of each mandible were aseptically extracted and any remaining extraneous soft tissue removed followed by removal of bone approximately 2 mm anterior and posterior to the three molar teeth. Each pair of molars were sonicated on ice in 1 ml sterile PBS, pH 7.4, in siliconized 2 ml microcentrifuge tubes, the molars aseptically removed and approximately 5 × 10^8^ depurinated cells of *S. mitis* UF2 then added, followed by vortexing for 5 sec and centrifugation (10,000 × *g*, 10 min at 4 °C). Cell pellets were then processed for DNA isolation as described for swabs. Recovered bacterial genomes in DNA samples were assessed by qPCR in 20 μl reactions run in triplicate in a Bio-Rad CFX96 real-time PCR instrument using 10 μl SsoAdvanced Universal SYBR®Green Supermix (Bio-Rad, Hercules, CA), 0.5 μl each primer and 9 μl of DNA diluted in 4 mM Tris-HCl, pH 8.0. Samples were run for 3 min at 98°C followed by either 34 cycles (*rpsL*) or 40 cycles (98°C, 15 s; annealing/elongation, 45 s at 70°C for *S. mutans*, 55°C for *rpsL* and 64.5°C for *S. sanguinis*); followed by a melt curve from 65-95°C at 0.5°C increments. Primers were as follows: *S. sanguinis* (Forward, 5’-GAGCGAATCATCAAGGATCAAAC-3’, Reverse, 5’-CGAGCAATAGCTTT-CGTAATAGG-3’), *S. mutans* (Forward, 5’-TGGCAAGTCCTGATGGTTTGAC-3’, Reverse, 5’-GGAAGCGGAAGCTGTGATGAAC-3’), *rpsL* (Forward, 5’-CCKAAYTCNGCNYTNCGT-AA-3’, Reverse, 5’-CGHACMCCWGGWARGTCYTT-3’). Primers were used at final concentrations of 0.50 μM (*S. mutans*), 0.15 μM (*S. sanguinis*) and 2.5 μM (rpsL). Standard curves were derived from DNA samples isolated from each strain grown to mid-exponential phase in BHI. *S. mutans* UA159 was used as standard for *rpsL* assays. Efficiencies, slopes and *r*^2^ values for standard curves were greater than 90%, −3.226 and 0.975, respectively. Results were analyzed using the Bio-Rad CFX Manager program. Under the listed conditions, all assays were specific for the targeted genomes. All work was performed in a BioSafety cabinet under aseptic conditions.

## Data analysis and presentation

Statistical tests were performed in GraphPad Prism 9.0 (graphpad.com) or R (Team, 2018). Significance was determined by statistical tests indicated in the figure legends. *P*-values ≤ 0.05 were considered significant. DESeq2 calculations of RNA-Seq datasets were completed in Geneious Prime 2021.1 or in the pcaExplorer R package (Marini & Binder, 2019). Confidence intervals (95%) of replicate samples were determined by the pcaExplorer R package.

## Acknowledgements

We would like to thank Seon-Sook An, Karina Kunka, Shannon Green, and Jody Turner for their advice and technical assistance.

## Author Contributions

TP and TK designed the *in vitro* experiments and DJC and RAB designed the *in vivo* experiment. TP completed the *in vitro* experiments and ACS, CAL and ASB completed the *in vivo* experiment. TP, DJC, RAB, and TK wrote the manuscript. All authors reviewed and approved the manuscript.

## Abbreviated Summary

*Streptococcus sanguinis* is an oral bacterium that competes with *Streptococcus mutans*, a causative agent of dental caries. We report that growth of *S. sanguinis* SK36 in acidic media leads to decreased manganese levels and changes in carbon catabolite repression. A knockout mutant of the primary manganese transporter grows poorly in acidic media but shows similar colonization and competition with *S. mutans in vivo* when compared to wild type.

## Supporting Information

Puccio_et_al_Supp_Tables.xlsx Excel document Supplementary Tables

Puccio_et_al_Suppinfo.pdf PDF document Supplementary Figures

