## Supplementary Material for "Manganese transport by *Streptococcus sanguinis* in acidic conditions and its impact on growth *in vitro* and *in vivo*"

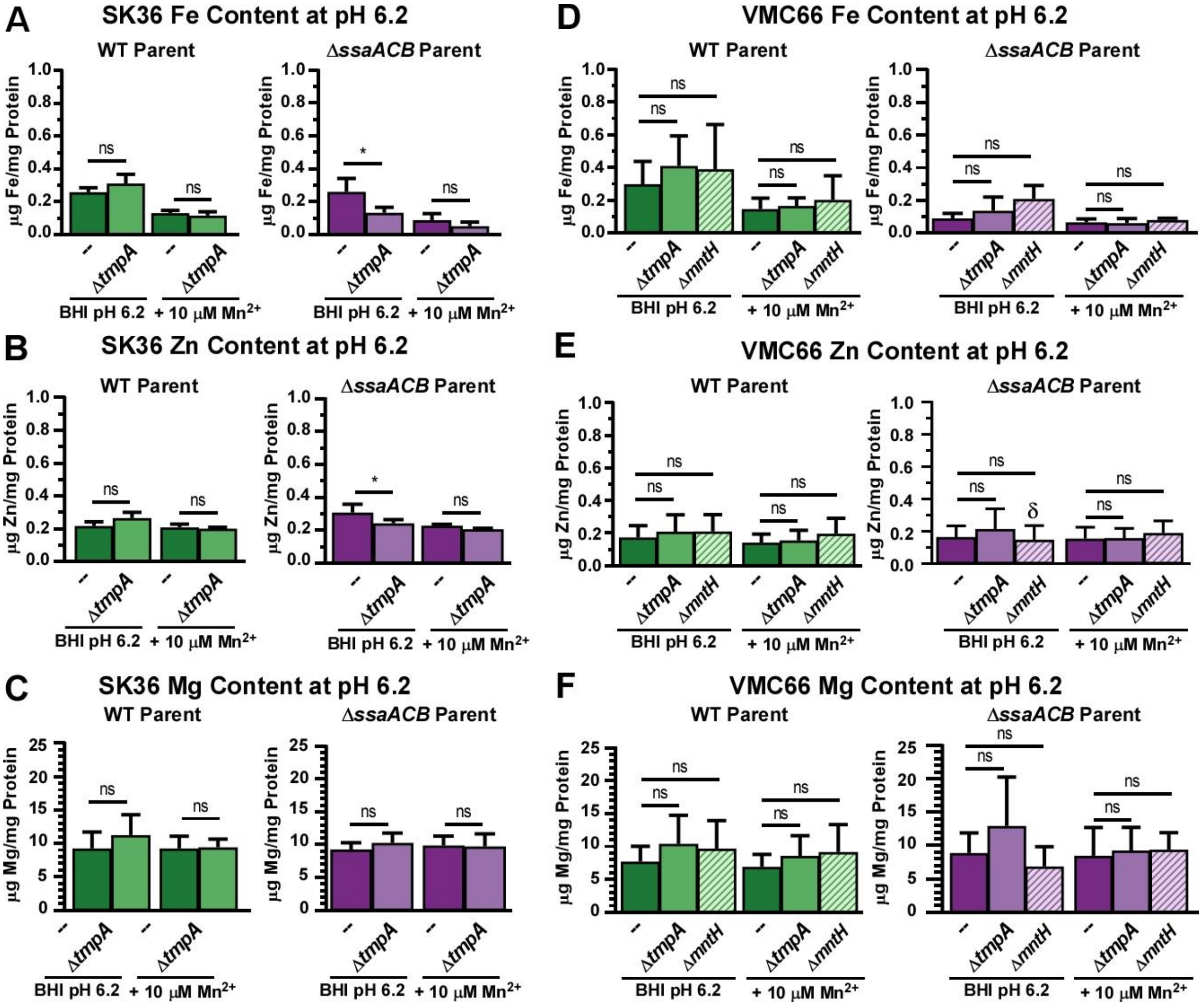


**Figure S1. Metal content of manganese-transporter mutants of SK36 and VMC66 in pH 6.2 BHI**

Impact of reduced pH on levels of iron (A & D), zinc (B & E), and magnesium (C & F) was assessed using ICP-OES on SK36 (A-C) and VMC66 (D-F) manganese-transporter mutants. Cells were grown in BHI at pH 6.2 ± 10 µM $Mn^{2+}$. Significance was determined by one-way ANOVA with Bonferonni's multiple comparisons test for each mutant as compared to the parent under the same conditions (* $P < 0.05$). Bars with δ had values that fell below the lowest standard.

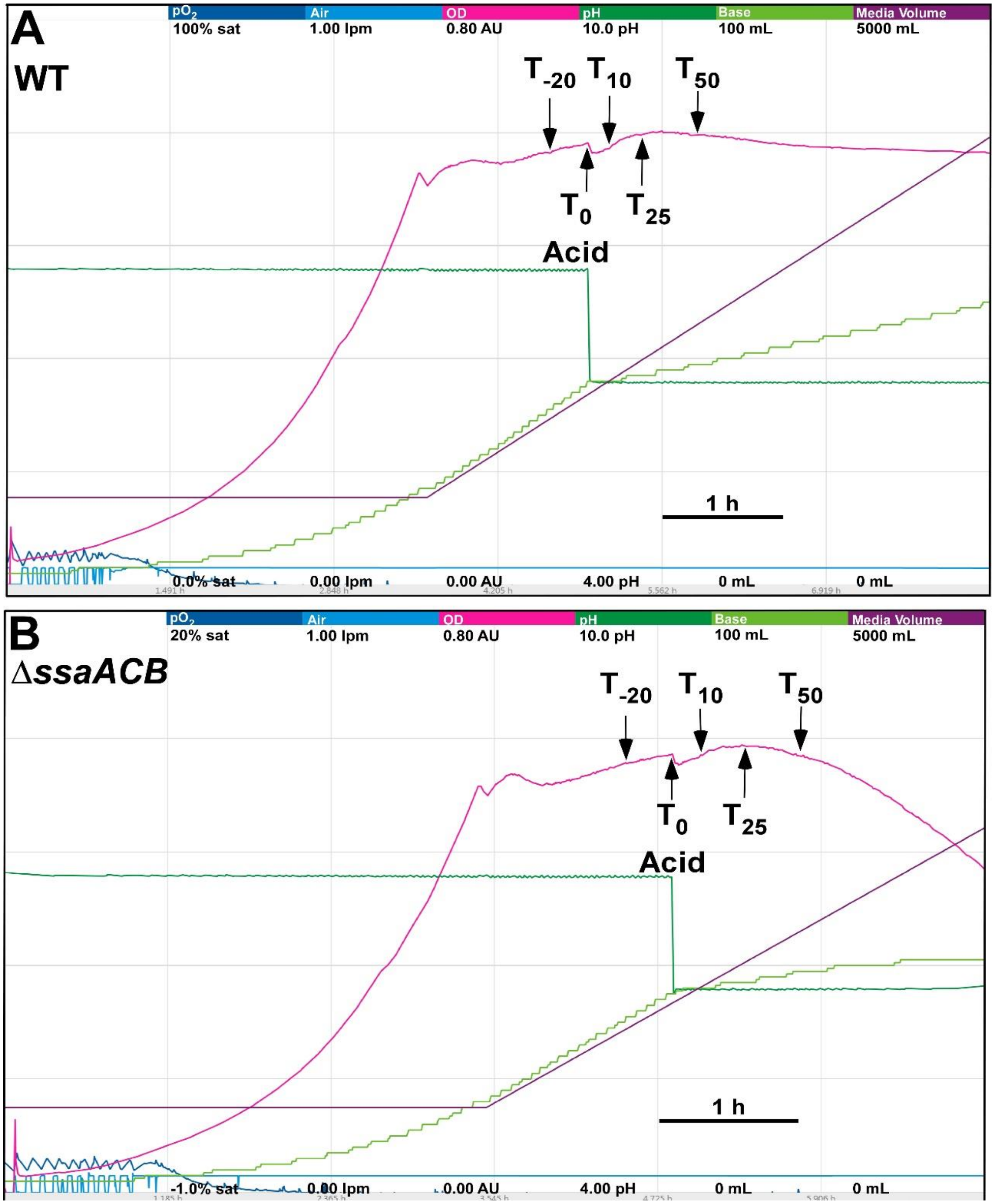


**Figure S2. Fermentor growth of WT and Δ*ssaACB* cells after pH reduction**

Representative charts for fermentor growth of *S. sanguinis* (A) WT and (B) Δ*ssaACB* cells. Each color represents a different parameter: cyan - air flow (liters per min; lpm), pink - optical density (840-910 nm; absorbance units; AU), dark green - pH, light green - base input (KOH), purple - media input (total volume). The scale for each parameter is indicated by the values under each respective parameter label (minimum at the bottom, maximum at the top). The time scale is indicated by the bar in the bottom right portion of each chart. Cells were grown under low oxygen conditions with pH reduced at 80 mins ($T_0$) after the media input and output pumps were turned on. Each sample time point is labeled.

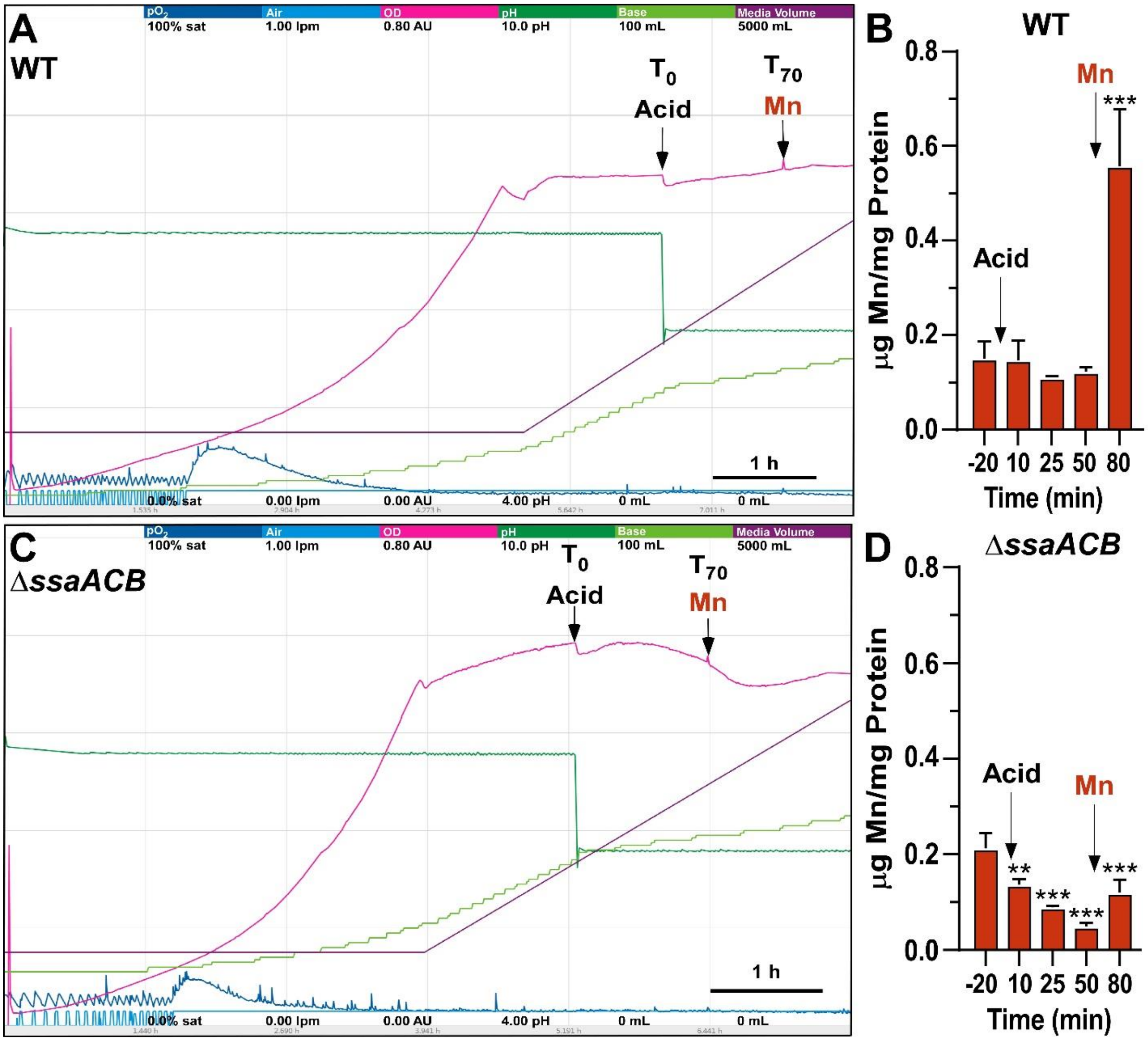


**Figure S3. Fermentor growth and metal content of WT and Δ*ssaACB* cells after pH reduction and addition of 10 µM $Mn^{2+}$**

Representative charts of three replicates of the (A) WT and (C) Δ*ssaACB* mutant with $Mn^{2+}$ added to 10 µM at 70 min post-pH reduction. Samples were collected for metal analysis at each previously described time point as well as $T_{80}$. Chart description is the same as in Figure S2. Manganese levels from fermentor cultures were measured for (B) WT and (D) Δ*ssaACB* using ICP-OES. Significance was determined using a one-way ANOVA with Dunnett's multiple comparisons tests where each pH 6.2 time point was compared to $T_{-20}$ (pH 7.4). $^{**}P < 0.01$, $^{***}P < 0.001$.

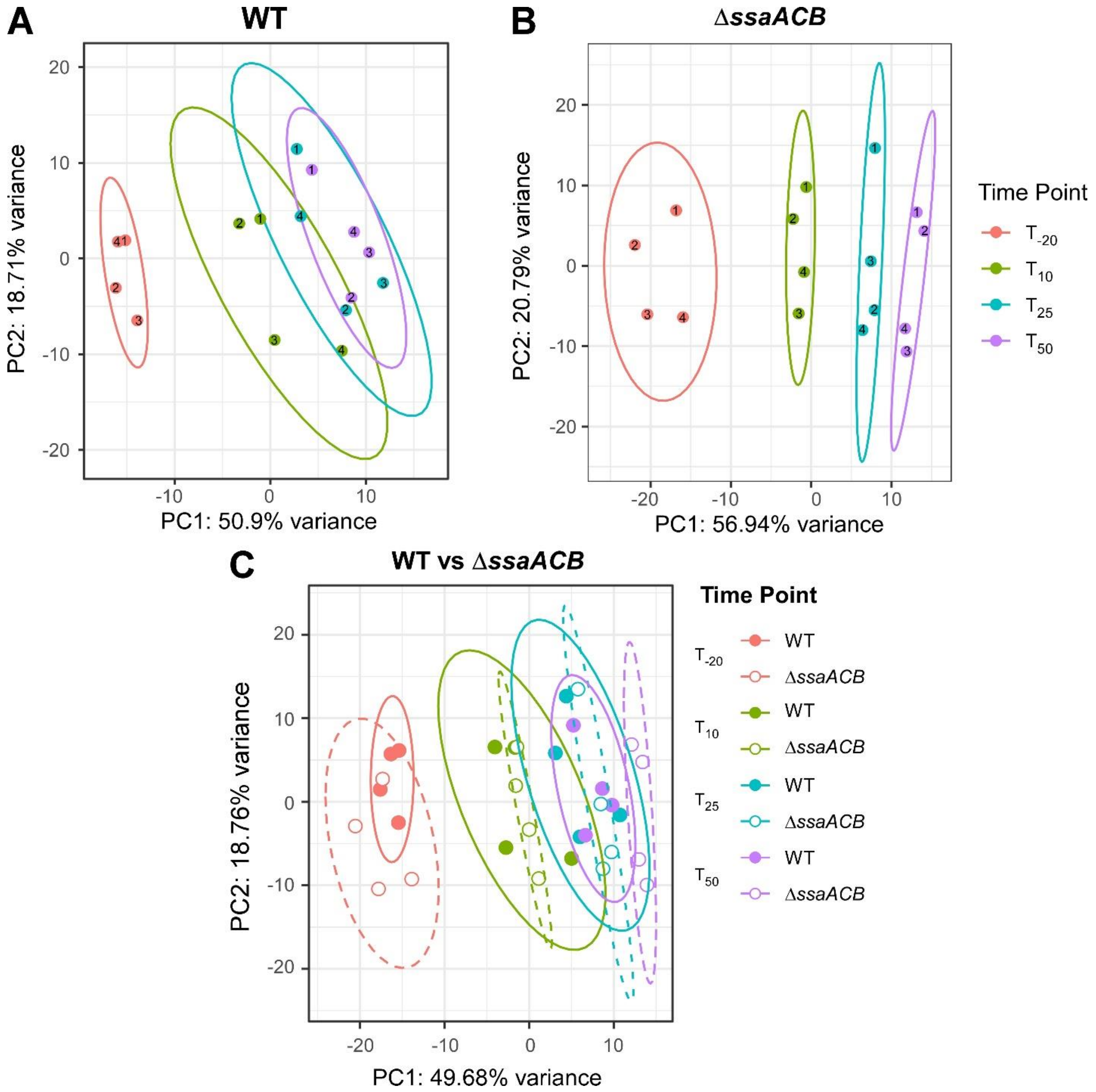


**Figure S4. Multivariate analysis of RNA-seq results**
PCA of RNA-seq results: (A) WT, (B) Δ*ssaACB*, and (C) both. Numbers in each circle in A and B represent the fermentor run number for that strain. In (C), filled circles and lines depict WT and empty circles and dashed lines represent Δ*ssaACB*. Ellipses represent 95% confidence intervals.

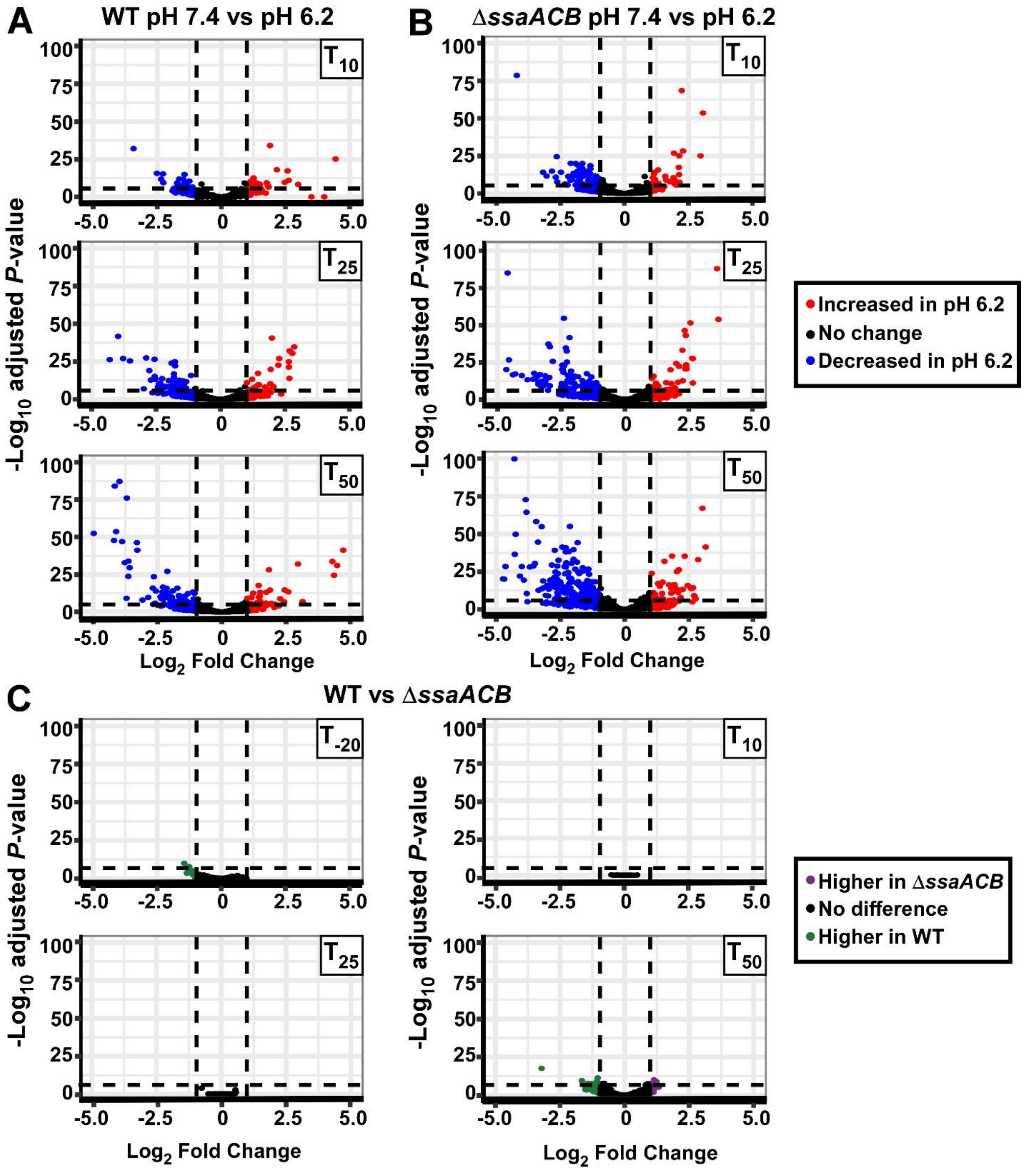


**Figure S5. Volcano plot representation of RNA-seq results**

Volcano plots comparing each pH 6.2 time point to $T_{-20}$ in WT (A) and Δ*ssaACB* (B) were generated using $\log_2$ fold changes generated in Geneious in the EnhancedVolcano package for R. Genes that are upregulated in the post-acid time point are positive on the x-axis (red) and those that are downregulated are negative (blue). Volcano plots comparing WT to Δ*ssaACB* at each time point (C). Genes that were more highly expressed in WT are green and those that were more highly expressed in Δ*ssaACB* are purple.

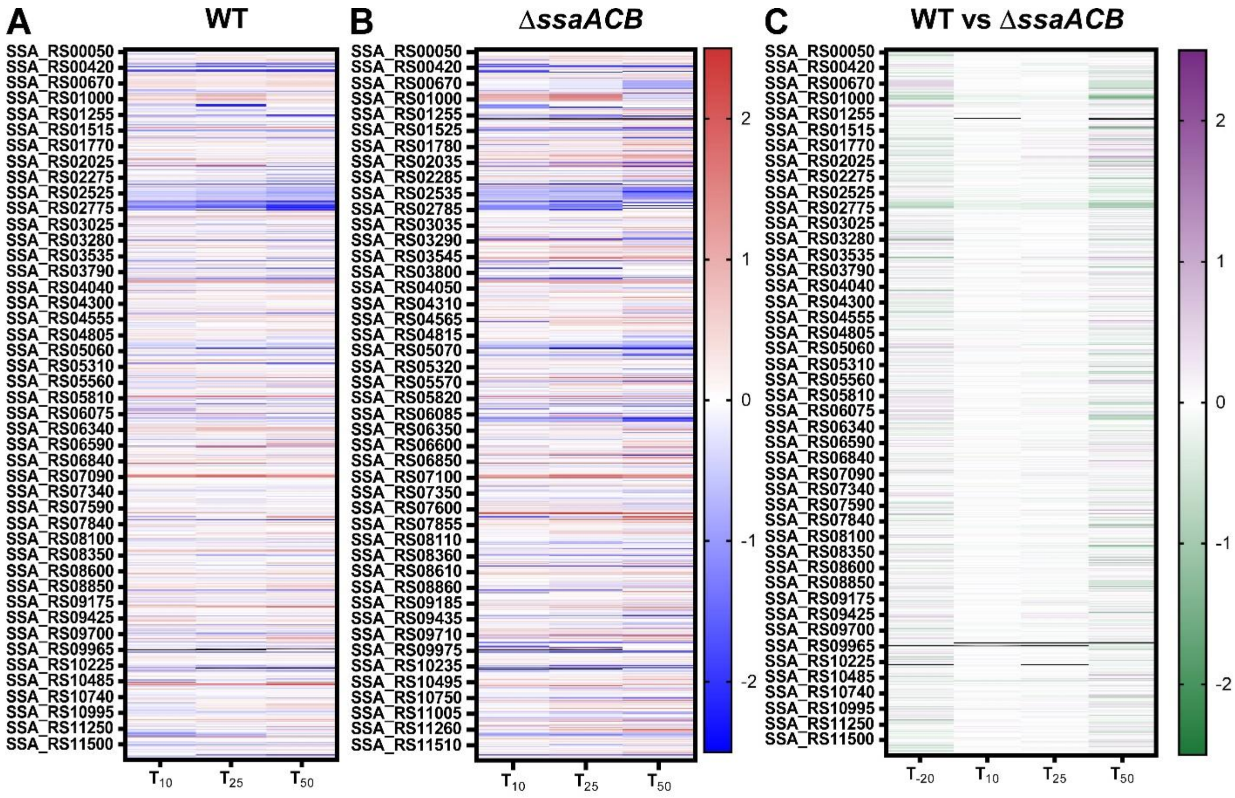


**Figure S6. Heatmaps of RNA-seq analysis**

Heatmap displaying the $log_2$ fold change values of each gene at the indicated time point as compared to $T_{-20}$ for WT (A) and Δ*ssaACB* (B) strains. Positive $log_2$ fold change values (red) are upregulated in later samples as compared to $T_{-20}$, while negative values (blue) are downregulated. (C) Heatmap comparing $log_2$ fold change values between the two strains at each time point where genes that were more highly expressed in WT are green and those that were more highly expressed in Δ*ssaACB* are purple.
